# Genome expansions and regulatory contact entanglement help preserve ancestral metazoan synteny

**DOI:** 10.64898/2026.04.14.717158

**Authors:** Yehor Tertyshnyk, Thea F. Rogers, Darrin T. Schultz, Satoshi Takenawa, Bibudha Parasar, Fatih Sarigol, Abidin Erdost Irmak, Lynn Wachelder, Florian Stallovits, Jiesi Gang, Koto Kon-Nanjo, Tetsuo Kon, Frédéric Bantignies, Longzhi Tan, Oleg Simakov

## Abstract

Chromosomes constitute deeply conserved evolutionary units in many metazoan genomes, with chromosomal fusions and fissions, accompanied by sub-chromosomal rearrangements, rewiring three dimensional genome architecture. How chromatin loops and compartments that define distal regulatory interactions within chromosomes impose functional constraints that affect this long-term evolutionary process (and vice-versa) is an emerging research topic. Genome expansions, especially through transposable element (TE) activity, test these constraints by increasing the genomic distances over which regulatory interactions must function and were thus suggested to be the drivers of chromosomal rearrangements. To study dynamics and stability of such distal interactions in the light of genome expansions, we focus on the cnidarian *Hydra vulgaris*, which, based on its simple and well-understood biology as well as one of the largest genomes among cnidarians, is particularly suited to test how chromatin loop and genome architecture respond to genome expansion. We investigate genome architecture using Micro-C, single-cell Hi-C (Dip-C), and DNA FISH, and perform comparative analysis using available genomic and epigenomic data. Contrary to prior expectations, our analysis of whole-genome data and particular loci (e.g., Wnt) suggests a scenario where genome expansion did not only result in chromatin loops often reaching several megabases in hydra, but also led to regulatory contact mixing and entanglement, introducing additional constraints to maintain ancestral genomic architecture. Generalizing these findings across hundreds of metazoan genomes, we show a new mechanistic role for genome expansion in yielding entangled long-range regulatory configurations that, in turn, decelerate chromosomal rearrangements, thus maintaining (and not breaking) ancestral regulatory states and synteny.

**Significance statement:** Some of the largest and most repetitive animal genomes retain a surprisingly high level of deeply conserved metazoan synteny. As repetitive regions are often associated with chromosomal rearrangements, it has been enigmatic why syntenic retention is so frequently observed. In this study, using hydra as a model system for both stem cell biology and an evolutionary history rich in transposable-element driven genome expansion, a multi-level conformational landscape dissection reveals a multitude of long-range regulatory states. We show that mixing of multiple such regulatory links accompanied by genomic expansion is associated with maintained ancestral synteny, thus pointing to the counter-intuitive role of genome expansions as “fossilization” agents across metazoan genomes.

## Introduction

What drives the conservation of the ancestral genome organization and what are the prerequisites for some species to undergo extensive genome rearrangements is an emerging topic with substantial implications for biodiversity research (1–3). Among many genomic features that have been studied in the past years (1), genome sizes were found to vary substantially across the animal kingdom (3): from the smallest genomes of free-living (i.e., non-symbiotic or -parasitic) animals of a few dozen megabasepairs (Mbp) to over a hundred gigabasepairs (Gbp) (4). Initial expectations were suggesting that accumulation of transposable elements (TEs) may facilitate hotspots of recombination, breaking genomic arrangements at intra- and, possibly, inter-chromosomal levels (5). On the contrary, some of the largest animal genomes retain a substantial part of ancestral synteny (6, 7), whereas smaller genomes are some of the most syntenically scrambled (8, 9). So far, no study has attempted to address this conundrum from both evolutionary and mechanistic perspectives. The central question whether genome size, either through TE accumulation or recombination-driven purging, can play a major role in shaping evolutionary trajectories is yet to be explored.

Our previous work has established the concept of irreversible fusion-with-mixing (FWM) characters at the chromosomal level, resulting in mixed chromosomal states that are retained as shared derived (synapomorphic) characters in evolution (10, 11). While such characters were found to clearly demarcate ancient animal clades (12), the functional implications of these chromosomal-scale events were unclear. In our past works (13, 14), we hypothesised that FWMs’ primary long-term evolutionary contribution is the generation of novel regulatory combinations, by bringing previously unlinked genes and their associated enhancers onto the same chromosome (**Figure 1**). More recently, we generalized this finding to sub-chromosomal level suggesting that mixing of enhancer-promoter contacts results in “entangled” regulatory configurations that constrain these loci against further translocations, locking in and shaping evolution of two or more otherwise separately regulated loci (15). In that view, FWM is happening at both chromosomal and sub-chromosomal levels, yet through different evolutionary constraints (meiotic on chromosomal (16–18) and regulatory at local linkage levels, respectively (14)).

**Figure 1.**
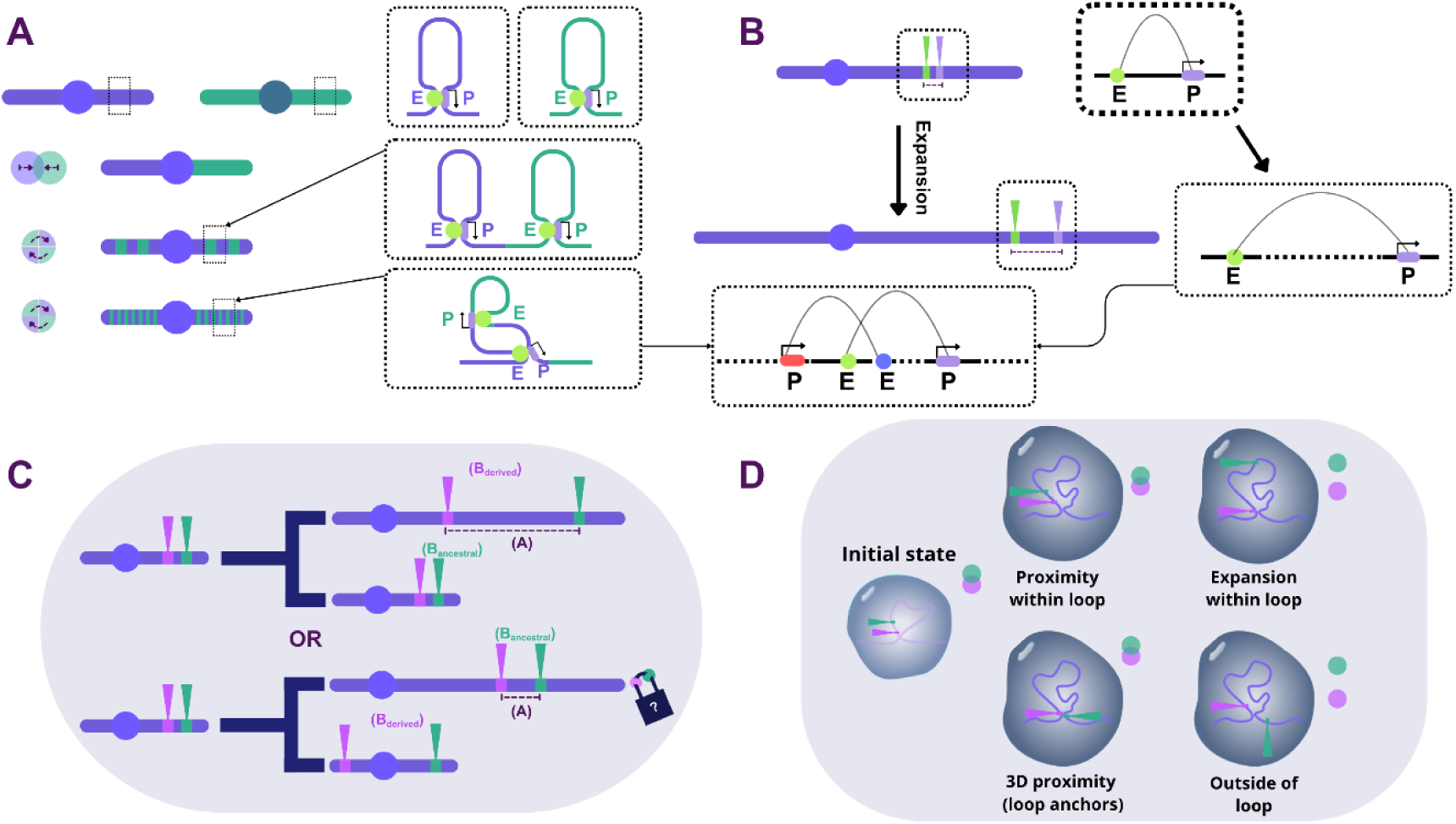
Hypothesis of regulatory mixing and entanglement and the role of genome expansions. (A) Fusion-with-mixing (FWM) may result in mixed (entangled) regulatory neighbourhoods consisting of multiple overlapping enhancer-promoter contacts. These configurations are predicted to be maintained in evolution as “disentanglement” via inversions may break functional regulatory linkages and origin of novel regulatory links within these loci will furthermore obscure the ancestrally separate states. (B) Genome expansion may impose an additional constraint on, or prolong enhancer-promoter entangled state persistence as more inversions may be required to disentangle the contacts. (C) Two hypothetical scenarios of the impact of genome expansion on loci proximity and which species would be considered more ancestral; genome expansion correlates with the distance and the cluster of two genes is maintained in the smaller genome (C:top) or the genome expansion enables persistence of a gene cluster retaining the ancestral state, whereas the gene cluster is broken in a smaller genome (C:bottom). (D) Different scenarios how genome dynamics may shape loci proximity in 3D space and thus the readout of Hi-C and DNA FISH experiments.

The key prediction of this hypothesis is that both the amount of mixed regulatory contacts, as well as the genomic distances that are spanned by them may constitute a genomic “balance” on how fast intra- and inter-chromosomal rearrangements can accumulate over the macro-evolutionary time-span. To test this hypothesis, phylogenetically key model systems where the effect of genome expansion could be studied in the context of regulatory contacts have been lacking. First, many invertebrate genomes are relatively small, secondly, little high resolution regulatory data (such as chromosomal conformational capture, e.g., Hi-C) is available (2, 19), and, thirdly, some systems that show both genome expansion and have established genomic resources are morphologically complex (e.g., cephalopods (13)).

In this study, we therefore aimed to investigate the role of genome expansion on emergence and maintenance of long-range contacts in a cnidarian *Hydra vulgaris*. Cnidaria (such as anthozoans, including corals, and hydrozoans, including hydra) are morphologically simple animals with two distinct epithelial layers (ecto- and endoderm) encompassing a gastric cavity. Despite this simplicity, cnidarians have emerged as key evolutionary model systems in numerous development and cell type evolution studies (20–24) due to the retention of ancient metazoan genomic and transcriptomic features (25). Among cnidarians, the hydrozoan *Hydra vulgaris* is particularly well suited to study the role of genome expansion through TEs in the evolution and maintenance of gene regulation (26, 27).

Firstly, hydra is one of the most well established cnidarians in the laboratory that has been a historical foundation for modern-day developmental biology, with the animal regeneration capability first described as well as organizer transplantation experiments conducted in this species (28–31). Hydra’s stem cell biology and several major stem cell lineages (e.g., ecto-, endo-dermal, and the interstitial, or i-cells, stem cell lineages) have been extensively studied (32) with implications for the species’ apparent lack of ageing (33, 34). More recently, cnidarians became emerging models for regulatory interaction evolution as they lack CTCF and topologically associating domain-like compartments (35, 36), yet as many eukaryotes they have cohesin and the associated loop extrusion (36–40). Despite these recent insights, since most cnidarian genomes are small, with high gene density, the regulatory interactions studied so far were limited to very proximal genomic regions (35). Furthermore, no clear long-range chromatin loops were observed (37) and no chromatin loops detected in Hi-C in basally branching metazoans have been validated as physical entities, e.g., via imaging. The second advantage of hydra, and brown hydras in particular (26, 41), to address this question is therefore their large (around 1Gbp) genome, one of the largest among cnidarians. This genome size was found to be a result of expansion activity of a diverse set of TEs, some of which are active in stem-cell lineage specific manner (26, 27, 41). For many years, it has been suggested (41) that this genome expansion, and the observed higher rates of protein sequence evolution (41), makes hydra a highly derived genome, yet more recent synteny analyses have shown a very well conserved chromosomal complement (11). The expanded genome of hydra, along with its molecular tools, thus enables us to test the hypothesis that chromatin loop size is a function of the genome size and test its impact on the maintenance or loss of synteny.

In this study, we conduct high-resolution chromatin conformational data (Micro-C) analysis of the hydra genome to unravel chromatin loop landscape and, together with several publicly available epigenetic datasets, characterize its regulatory features. Compared to the standard conformational capture (Hi-C) protocols, Micro-C enables detection of chromatin loops at much finer (nucleosome-level) resolution. These analyses enable us to identify very long-range interactions in this species, such as 4Mb+ chromatin loops around the Wnt ligand cluster. We validate these conformational data predictions by DNA fluorescence *in situ* hybridization (DNA FISH) on intact nuclei and assess its variability with single-cell Hi-C protocol (42). Through comparative genomic analysis, we find that chromatin loops in hydra are correlated with ancestral syntenic regions, suggesting that the genome expansion leads to less macro-evolutionary rearrangements within animal chromosomes. Through novel comparative genomic approaches to estimate genomic mixing among hundreds of animal genomes we can generalize these findings to many other metazoan clades. Our study thus provides a mechanistic explanation of the proposed regulatory entanglement in animal genomes and reveals the role of genome expansion as a genomic “fossilization” agent enabling preservation of deeply conserved synteny at both chromosomal and sub-chromosomal levels.

## Results

### Micro-C reveals the presence of long-range chromatin loops in a cnidarian

The hydra genome maintains ancestral syntenic configurations as represented by the retention of complete metazoan ancestral linkage groups (11, 27), however the maintenance of linkages at local sub-chromosomal scale and its regulatory landscape has not been extensively studied so far. We applied a micrococcal nuclease conformational capture (Micro-C) approach that enables high level resolution of conformational contacts. This allowed us to investigate the chromatin loop presence and their properties in the context of the expanded genome of hydra. While Hi-C approaches have generally been applied to non-bilaterians (35, 37), the smaller genomes used in these studies as well as lack in the resolution or the reference assembly quality did not permit observation of long-range interactions.

Bulk whole-body Micro-C data analysis supports the presence of Rabl-like chromosomal configuration (**Figure 2a**, **Figure S1,7**), previously reported and linked to Condensin II loss (36). In such an arrangement chromosomes retain U-shape, with centromeres and telomeres being spatially separated. Moreover, similar to the previous studies (35–37), our Micro-C data shows no evidence of topologically associating domain (TAD)-like structures, which is corroborated by the CTCF absence in non-bilaterian genomes (35). Interestingly, our data clearly reveals presence of long-range (1 Mb and above) chromatin loops in hydra (**Figure 2a,d**), which has been elusive in previous studies on basally branching metazoans, where median loop sizes were typically within the range of tens of kilobases only (only a single 1 Mb loop in *Nematosella vectensis* was identified, 37). Overall we detect 544 loops (union between 316 detected loops by Chromsight (43) and 228 by Mustache (44)). The median size of the detected loops was 2.2 Mb (mean 3.34 Mb). Similar to sponges (37), inspection of Micro-C maps revealed symmetrical jets (45), often on loop anchors, a signature of dynamic loop extrusion in CTCF absence (45, 46) (**Figure S1e,f**). Around 60% of loop anchors had open chromatin (ATAC-seq) peaks, with almost 30-fold enrichment compared to the rest of the genome (**Figure S2**). Motif enrichment analysis was conducted for loop anchors, revealing a de-novo motif significantly enriched in loop anchors with 25% of the loops having it in at least one of the two anchor bins (**Figure 2**h-i). The mechanism of loop extrusion and control across animals is far from understood. We searched the hydra genome for several reported components and found presence of PDS5, which has been recently reported as a key component in active loop extrusion (44), suggesting possibly deep evolutionary conservation of its function.

**Figure 2:**
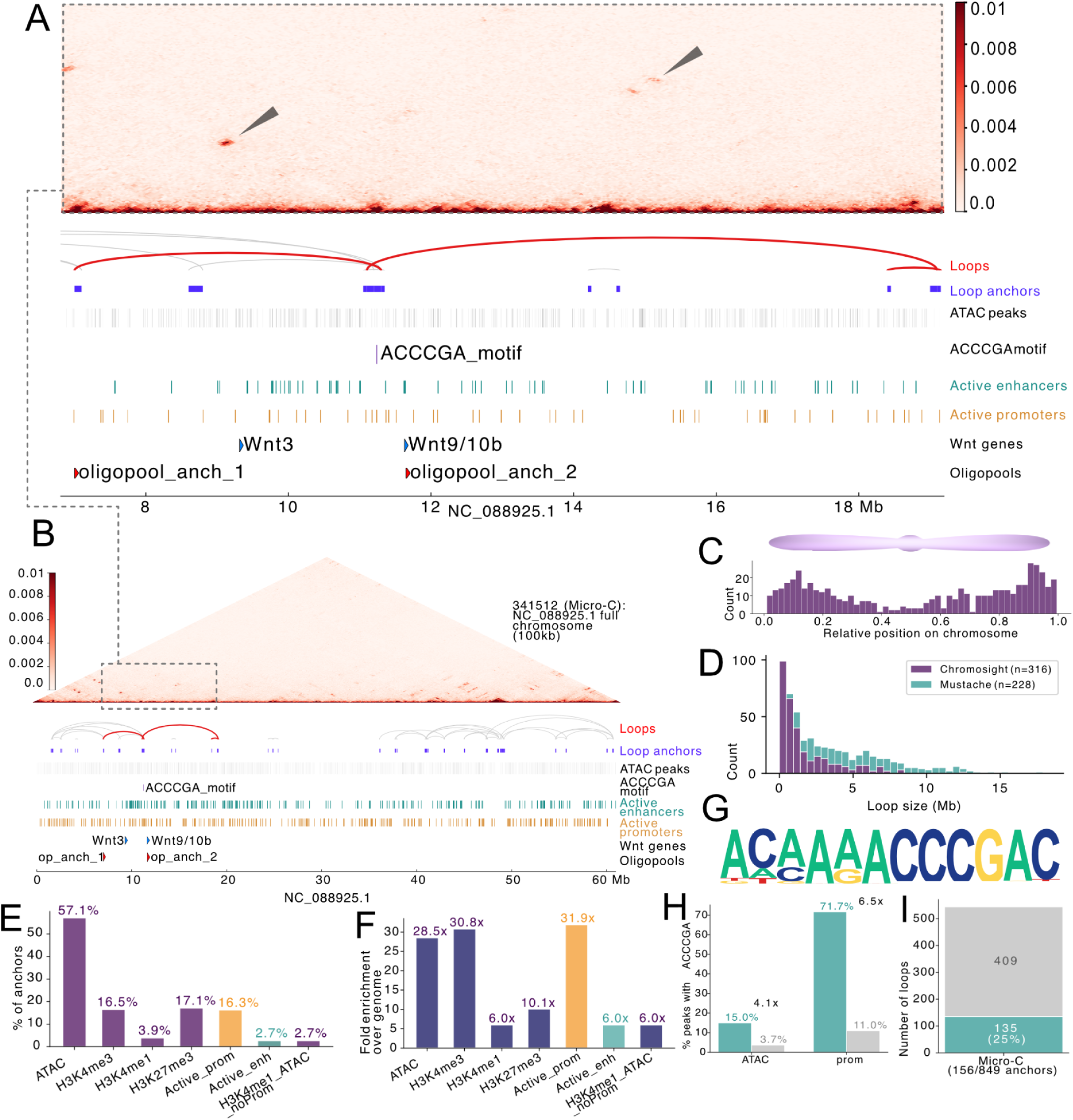
Micro-C reveals long-range interactions in the expanded genome of hydra. (A) Zoom-in to region 6.80-19.1Mb showing two “entangled” loops (red) containing Wnt3 and Wnt9/10a,b, as well as ATAC-seq peaks, enhancer and promoter calls. (B) Micro-C map of Chromosome 6 (NC_088925.1) at 100 kb resolution showing enrichment of nested loops on both ends of chromosome and region of interest. (C-I) Loop calling, statistics and motif enrichment analysis; (C) Distribution of relative positions of loops across all 15 chromosomes of hydra. (D) Size distribution of *H. vulgaris* loops called with two different tools (Chromsight and Mustache), additionally central tendencies of loop sizes. (E) Fraction of loops containing ATAC seq peaks and other epigenetic marks. (F) Relative enrichment of epigenetic marks in loops vs. the genome-wide background. (G) De-novo discovered motif enriched in ATAC peaks found in loops. (H) Enrichment of the motif in (G) (and its reverse complement) at loop anchors (green) and genome wide (gray). (I) Fraction of loops having the motif in (G) at least one anchor (green).

Chromatin loops are not randomly distributed along hydra chromosomes. We find a trend of higher loop number on both ends of chromosomes and depletion in the centromeric region (**Figure 2c**), suggesting a potential regulatory impact of Rabl-like conformation in the hydra genome. Strikingly, we found that 60% of the hydra genome is covered by the predicted chromatin loops, most of which (65%) are nested inside at least one another loop.

To complement loop detection, we ran enhancer-promoter detection with Activity-by-Contact (ABC) (47) using available open chromatin and histone mark data for hydra (48), as well as our Micro-C data (Methods). This detected 40,785 putative enhancer-promoter (E-P) links (ABC score threshold of 0.05, Methods), with an average interaction distance of 0.37 Mbp and the longest E-P prediction of 5 Mbp. We then combined this data with phyloP conservation score analysis based on alignment of 10 cnidarian species and the chordate amphioxus (Methods). ABC-predicted enhancers showed moderate conservation compared to whole genome background with the mean phyloP of 0.207. Contrary to the detected loops, enhancer-promoter links were more evenly distributed along the chromosomes.

Overall, these results suggest the presence of a complex distal regulatory landscape in hydra, providing for complementary data to findings reported for a few select model organisms (49–51). Furthermore, our data shows presence of long-range interactions and often positionally overlapping chromatin loops that have not been observed before in other, smaller, cnidarian and non-bilaterian genomes (36).

### Chromatin loop landscape around known micro-syntenic linkages

To further assess the role of this topological landscape onto gene regulation we focused on known microsyntenic clusters. Many local gene linkages, such as Hox or Wnt have been reported (52–54) across metazoans and various regulatory models were proposed to explain their linkages (55–58). However, with very few exceptions such as that of the Hox genes (52), the functionality and co-regulatory aspects, if any, of these linkages are unclear. In this study, we focus on the Wnt clusters (59), as their presence has been widely reported in the literature across metazoans, yet no clear functional reason for their clustering has been determined so far. We profiled the Wnt complement across several metazoan species (**Figure S3**), which highlighted that despite general deep conservation of Wnt ligands across animals, their chromosomal locations and associations are dynamic in evolution. This makes the Wnt cluster particularly useful for studying how regulatory contact mixing may impact its evolutionary stability. We focussed on the Wnt3 and Wnt9/10a,b syntenic cluster in hydra (Wnt3-9/10), which, together with the Wnt5-7 linkage (**Figure S3**), is the major Wnt gene pair cluster across metazoans. Our Micro-C data in hydra revealed several chromatin loops forming a highly interlinked landscape in the region of Wnt3 and Wnt9/10a,b location. Specifically, we observed two large and overlapping loops (4.5Mb and 8Mb) around this locus in hydra: the 4.5Mb loop encompasses Wnt3 and one of its anchors localized closely to Wnt9/10a,b, which itself lies within the 8Mb loop span (**Figure 2A**).

The loop structure around the Wnt locus seems to be conserved in evolution. Using publicly available Hi-C data for the green hydra *H. viridissima* which diverged about 80 million years ago (26) from *H. vulgaris*, we were able to capture similar topological organization around the Wnt locus spanning roughly 2.5Mb (compared to approximately 10Mb of entangled loops around the Wnt3-Wnt9/10 locus in *H. vulgaris*, **Figure S5**), suggesting its evolutionary persistence in hydrozoans. Importantly, as there are dozens or even hundreds of genes that can be spanned by such long-distance loops, the effects on gene expression can affect multiple genes throughout the loop interaction distances (15).

To cross-validate loop predictions, we have analyzed the E-P predictions by ABC and the evolutionary conservation of the putative enhancers around Wnt loci. Compared to other E-P contacts, we observed an even higher phyloP score for all Wnt-associated ABC enhancers (0.383, n=26), with averages of maximum values within enhancer coordinates being 0.949 for WNT enhancers versus 0.645 for all enhancers. We then investigated the region on chromosome 6 containing Wnt3, Wnt9/10a,b loci and two overlapping loops (position 7 to 19Mb). This region showed to be densely interconnected with 563 ABC-predicted enhancer-promoter interactions. Five enhancers were predicted to regulate Wnt9/10a, and while no Wnt3 enhancer was detected at the ABC 0.05 threshold, three candidate enhancers were found at a more permissive 0.02 threshold. In addition to these direct E-P interactions, we identified 12 and 21 E-P links that span Wnt3 and Wnt9/10a,b (**Figure S4**), respectively, with the average length of 2.87 Mbp for Wnt3 and 2.61 Mbp for Wnt9/10a,b. This data reveals a high level of interconnectivity of this region and thus potential regulatory entanglement.

Together, topological analysis of specific loci shows evidence for multiple overlapping (mixed) regulatory contacts along hydra chromosomes, encompassing loci that are known metazoan microsyntenic linkages. These putative entangled configurations appear to be conserved in evolution at both conformational and enhancer conservation levels.

### Single-cell HiC shows dynamicity of regulatory loops across cells within the i-cell lineage

Chromatin loops are known to be very dynamic during development and across tissues and cell types (40, 60). In hydra, the three stem cell lineages produce a diversity of cell types and states (61). To complement Micro-C data which gives an average picture across all homeostatic hydra cells, we sought to investigate variability in chromatin loops in the multipotent i-cells at the single-cell resolution. The focus on the i-cell lineage is particularly informative as compared to the other stem cell lineages (ecto- and endo-dermal stem cell lineage) it gives rise to a variety of cell types, including gametes, neurons, or nematocysts (32) (**Figure 3**a). For this we isolated GFP-positive i-cells from the Cnnos+ transgenic *Hydra* vulgaris (62) using fluorescence-activated cell sorting (FACS), and conducted single-cell Dip-C library preparation and sequencing (42, 63).

**Figure 3.**
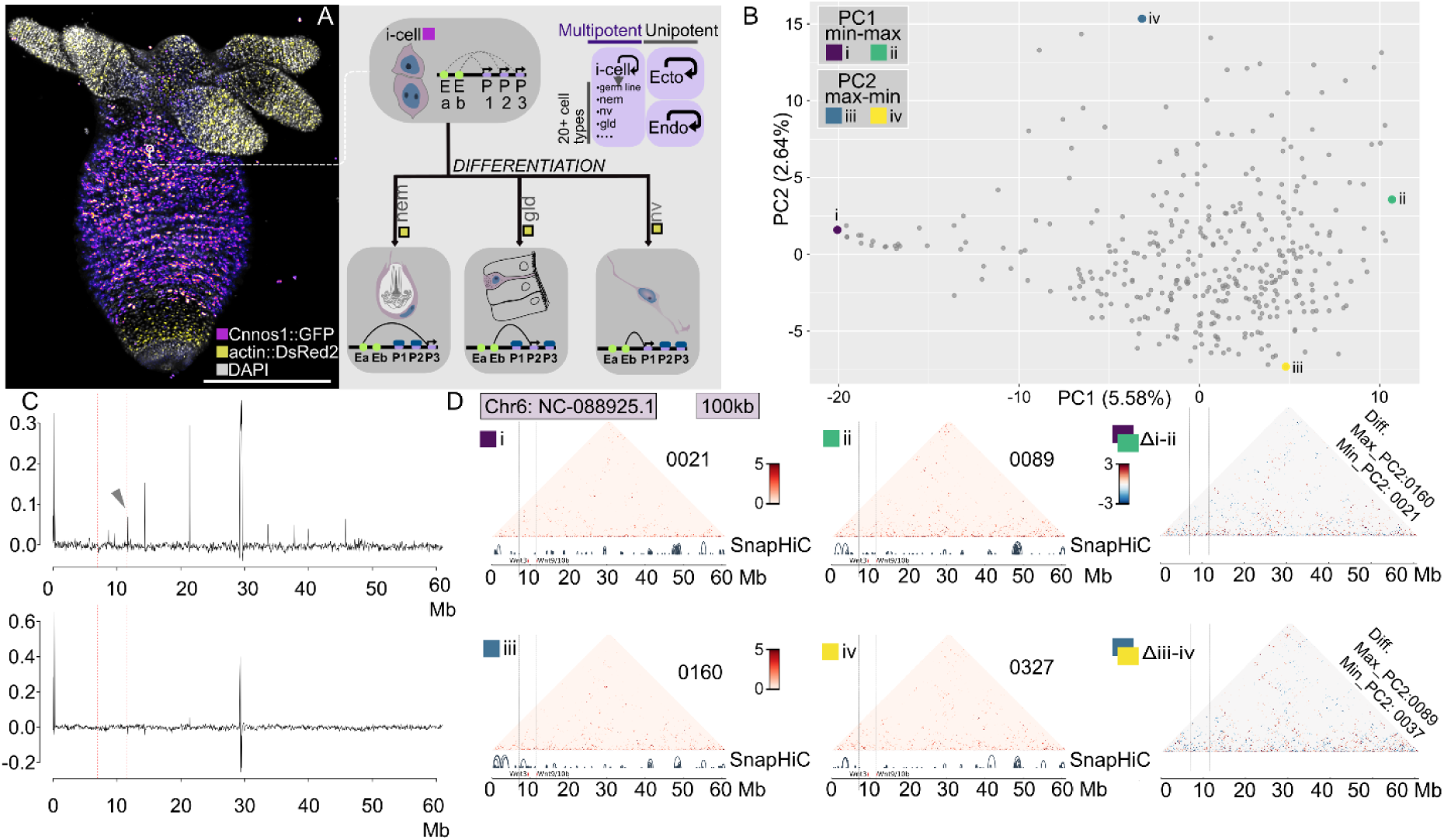
Dynamic and cell-specific contacts in the i-cell lineage revealed by single-cell Hi-C data. (A) Hydra i-cell lineage giving rise to several states and the expected different contact states within them. (B) Principal component analysis (PCA) of Dip-C pairwise contact intensity data (normalized contact intensity between 50kb bins) showing diversity and clustering of contact data, each dot represents a single cell. Examples shown in panel D are labelled as i, ii, iii, iv. (C) Contribution (loadings) of the first two principal components along the chromosome containing the Wnt3-9/10 cluster. Wnt cluster loop anchors are marked by red vertical lines. Variation along the PC1 axis contributes to one of the Wnt cluster anchors. (D) sc-HiC map examples for i, ii, iii, and iv showing Hi-C matrices for single cells, as well as SnapHiC loop calls at z > 1.96 threshold. Numbers for each Hi-C map indicate cell identifier. Hi-C difference maps between i and ii (PC1 extremes), and iii and iv (PC2 extremes) are shown in the final column.

Dip-C protocol has allowed individual profiling of 376 cells constituting the i-cell lineage (Methods). The data allowed us to visualize the dynamicity of topological structures within single cells (**Figure S6**). Dip-C matrices were largely consistent with Micro-C results, as evidenced by loop prediction done on “pseudo-bulk” (merged) Dip-C data (**Figure S2**). Focusing on chromosome 6, which is the location of the Wnt3-9/10 cluster, confirms the presence of the large loops visible in the Micro-C data (**Figure 3**, **Figure S6**). This data suggests that these chromatin loops are present in the i-cell lineage. Conducting loop calling on the single-cell data with the limit of 6Mb to loop size identified 26,326 loops on chromosome 6 across all 376 profiled cells (on average around 70 loops in each cell on chromosome 6 with the average length 0.62 Mb. Among these, 39 cells had loops around the Wnt9/10a,b, 41 cells had loops around Wnt3, and 7 cells had loops spanning both Wnt3 and Wnt9/10a,b, highlighting cell-type specificity and dynamicity of chromatin loop contacts within a single stem-cell lineage (64).

Clustering of normalized pairwise interaction values with principal component analysis (PCA) reveals a diversity of cells with distinct topological confirmations within the i-cell lineage (**Figure 3**b). This analysis identifies distinct regimes and chromatin loops emerging within this cluster that contributes to the separation of cells on the principal component space. Principal component (PC) 1 has a particular contribution around one of the loop anchors in the Wnt3-9/10 cluster (**Figure 3**c), suggesting dynamic, likely extrusion-based chromatin loop formation around these genes, playing an important role during i-cell development and differentiation. Principal component 2, on the other hand, shows contribution around centromeric and telomeric regions, suggesting less of a gene regulatory and a general chromosomal conformational role and a likely involvement in the Rabl configuration. This diversity of conformations is also reflected when inspecting individual cells representative of PC1 and PC2 extreme states (**Figure 3**d).

Together, both bulk and single-cell Hi-C data suggest presence of long-range chromatin loop interactions in the hydra genome. Their appearance ranges from single loops spanning few Mbp (e.g., **Figure S1**e) to hub-like structures, similar to the reported enhancer-promoter or promoter hubs (37, 65), but over a much larger (Mbp large) genomic scale (**Figure S1**b,c). The exact mechanism of this loop formation remains to be investigated. While distinct contact signatures (e..g, symmetrical jets) point towards specific cohesin-mediated loop extrusion (e.g., **Figure S1**f), which offers cell-specific contacts and regulatory landscape between open chromatin regions observed in our data. Equally, appearance of some hubs together with absence of cohesin extrusion signature, may also point to the presence of large (∼10 Mbp) polycomb-mediated repressive loop bodies (66–68) in hydra.

### Dynamic chromatin loop formations visualized by DNA FISH

While both Micro-C and Dip-C provide unique insights into the chromatin loop formation and their genomic properties in the hydra genome, current studies often lack the visual validation of such dynamics. We have applied a modified DNA fluorescence in-situ hybridization protocol (DNA FISH) to visualize interactions in intact hydra nuclei. Major modifications were required due to the AT-richness of the hydra genome and several digestion, incubation and probe design conditions were analyzed (Methods).

Using a 25mer highly abundant in centromeric regions (7693 copies in genome, 98.4% of which in centromeric regions, CP, **Figure S7**) and telomeric repeat we performed DNA FISH and investigated localization of centromeres (CE) and telomeres (TL). We found that 53 out of 85 (∼62%) nuclei exhibited CE/TL separation **(Figure 4a, Figure S7**) consistent with Rabl-like conformation as predicted by Micro-C (**Figure 4**a). This chromosomal arrangement is thus the prevalent type in hydra cells. The “mixed” cells with no clear CE/TL separation (32 out of 85, 38%) had lower median volume than the CE/TL separated cells (214.23 vs. 260.66 µm^3) and were more abundant among cells in the smallest volume quartile (103–178 µm^3; 63.6%). Although interphase Rabl-like conformation has been associated with the absence of Condensin II across eukaryotes (69, 70), it has also been shown to be cell-type and stage specific (71, 72). Cells at early developmental stages in *Drosophila*, *Xenopus* and human display Rabl conformation that is lost in further development suggesting its role in totipotency. This chromosome arrangement may be favored by rapidly dividing cells because it requires less reorganisation before mitosis than established chromosome territories (70, 71). Given that adult polyp of hydra is largely composed of lineage restricted stem cells (32) one can argue about the role of Rabl conformation in facilitation of rapid cell turnover in hydra and thus its association with stem cell identity. This is further supported by the observation that stem cell nuclei are larger (73) as well as a clear signature of Rabl conformation in our i-cell scHi-C data (**Figure S1,7**).

**Figure 4.**
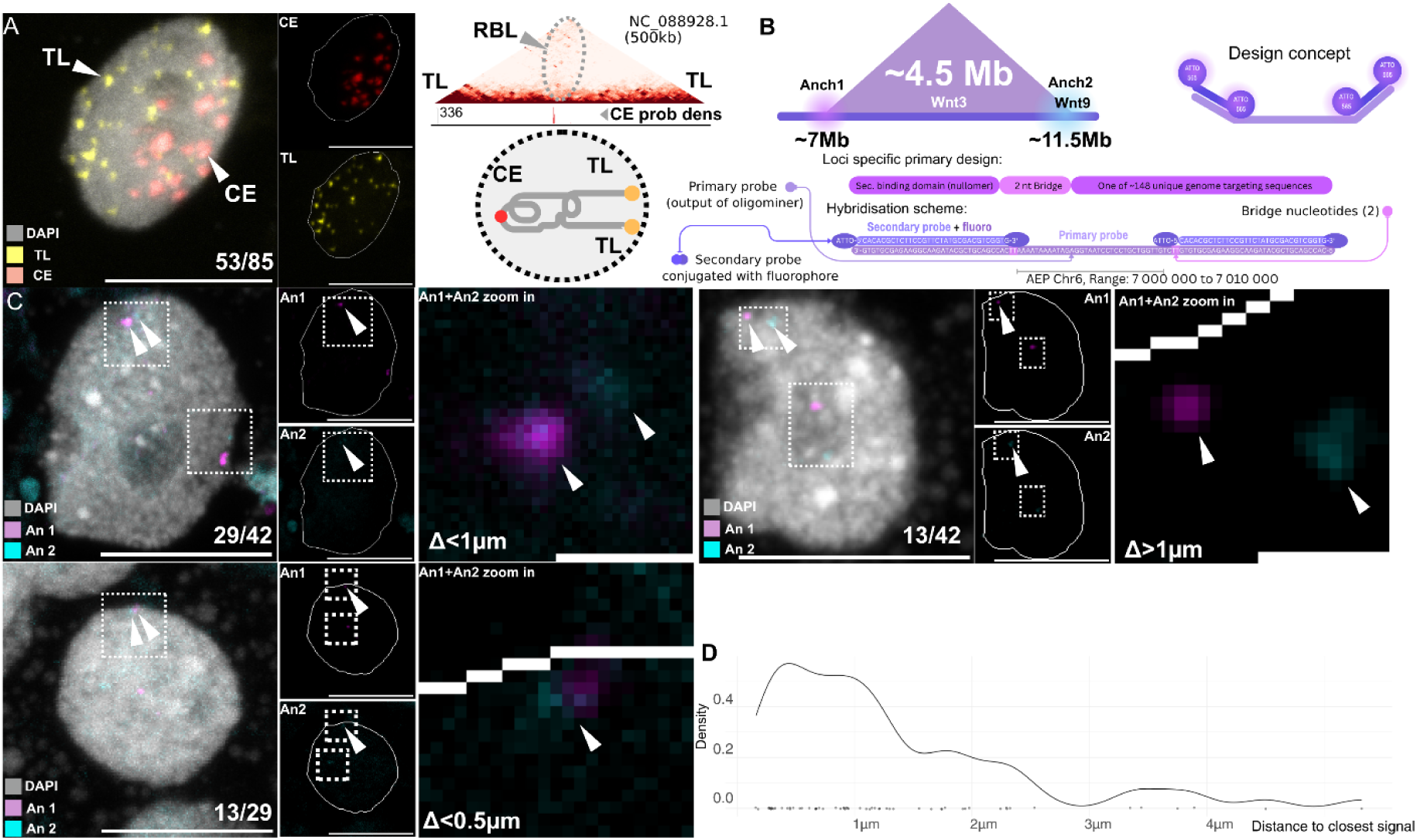
Compartmentalized chromosomes and dynamic contacts visualized in hydra cells. (A) Rabl-like conformation cross validated via Micro-C and with telomeric and centromeric DNA FISH. (B) Oligopool probe design for DNA FISH. (C) Different contact states of the Wnt loop observed for the Wnt cluster probes, as well as numbers of unique nuclei with minimal distances <1µm, <0.5µm and >1µm. Scales are 10µm and 1µm (zoom-in). (D) Distribution of distances between nearest DNA FISH signals.

Next, we designed oligopool probes against two regions separated by 4.5 Mb around the predicted Wnt cluster loop (**Figure 4b**). In more than half of the profiled nuclei (29/42, 69%) we found proximity or co-localization of <1 µm distance in at least one of the haplotypes, with around half of these anchor pairs (13/29, 45%, or 31% of the total 42) localized within 0.5 µm distance (**Figure 4c,d**). In another half of the profiled localizations, the loop anchors were farther apart than 1 µm and usually within the 2 µm range of each other. This second population points to a different type of anchor localizations likely reflecting random positioning within the nucleus.

Together with the single-cell Hi-C data, this visual validation highlights a very dynamic property of long-range chromatin loops in hydra. The nature of long-range chromatin loops suggests that multiple regulatory contacts within these regions can arise. Validation of the presence of these dynamic and specific contacts provides crucial additional evidence for the underlying functional mechanism behind the observed complex regulatory landscape (i.e., several, partially overlapping, chromatin loops) around the Wnt cluster.

### The expanded genome of hydra and maintenance of ancestral metazoan synteny

We next sought to investigate whether the presence of such long-range, developmentally dynamic and overlapping regulatory contacts along the chromosomes is reflected in syntenic organization of hydra and other metazoan genomes. A key prediction of the fusion-with-mixing mechanics of regulatory contacts is the emergence of regulatory constrained states and thus inhibition of exploration of chromosomal configurational space in evolution (“entanglement”) (14, 74). We have investigated the evolutionary impact using both simulation and comparative genomic approaches.

First, we have conducted simple inversion simulations for regulatorily separate and mixed states and found that especially in expanded and mixed enhancer-promoter contacts, the genes, even if functionally unrelated, tend to remain more closely positioned in the genomes (**Figure S8**). In our simulations, enhancer-promoter links that were overlapping took more generations to separate by chance. In case of expanded distances, such separation rarely took place during the complete length of the simulation.

Taking this prediction, we investigated empirical genomic data for the Wnt3-9/10 linkage and found that the genomic distance difference in anthozoans compared to hydrozoans was striking. In the expanded brown hydra genomes (such as *Hydra vulgaris*), the observed distance was 2.3 Mb, and smaller genomes of *Hydractinia symbiolongicarpus*, *Rhopilema esculentum*, and *Hydra viridissima* showed distances of 3.5 Mb, 1.4 Mb, and 0.5 Mb, respectively. In anthozoans, on the other hand, the genes of this Wnt cluster were spread across the chromosome or were lost: *Scolanthus callimorphus* and *Nematostella vectensis* showed around 12 Mb distance between Wnt3 and Wnt10, respectively (**Figure 5**a,b). Interestingly, the Wnt3-9/10 cluster was more tightly packed in bilaterians, with median distances of 0.36 Mb. Even in the highly expanded genomes of salamanders (*Ambystoma mexicanum*, 32 Gbp, and *Pleurodeles waltl*, 20 Gbp) the median region size of this Wnt locus was around 0.5 Mb. Our conformational data of multiple and dynamic chromatin loops around the Wnt cluster provides for the mechanistic explanation why such cluster has higher chance of persisting in evolution in its expanded state as suggested by the simulation (**Figure S8**).

**Figure 5:**
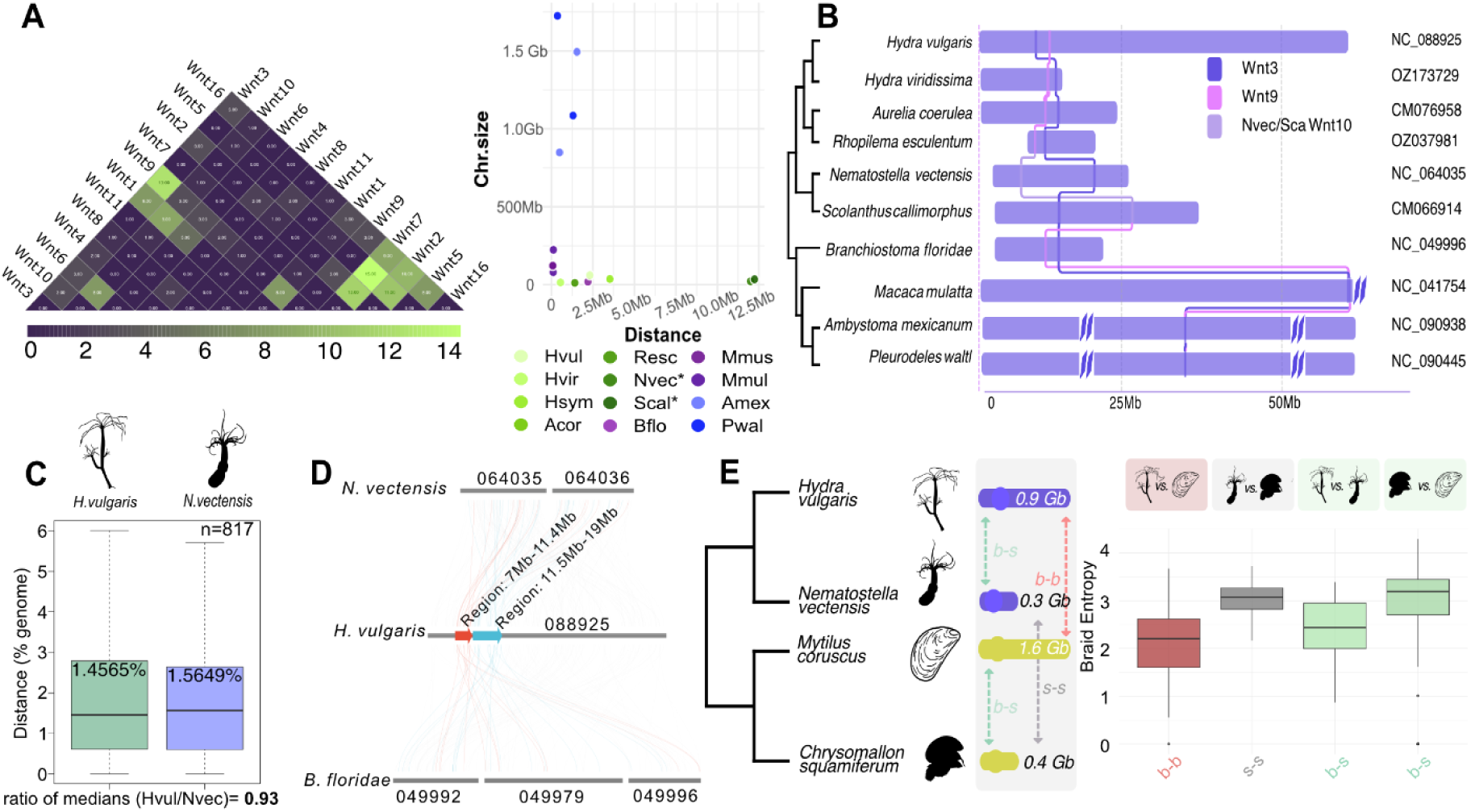
The effect of the genome size on gene linkage scaling and mixing. (A) Pairwise Wnt ligand distances and scaling across metazoans. Wnt3-9/10 linkage is constrained in many species, especially in species with very large genomes (e.g., axolotl) but is scattered in some species with small genomes (e.g., *Nematostella*). (B) Ribbon plot of clusters (Wnt) in expanded genomes, showing more constrained associations in larger genomes. (C) General orthology analysis shows that ortholog pairs that are constrained in bilaterians to a median distance of up to 2.5Mb are slightly more closely positioned in the expanded genome of hydra compared to *Nematostella*. (D) To assess the role of genome size in mixing within chromosomes we quantify the degree of positional shuffling between pairs of species. (E,F) This mixing analysis of braiding entropy of ortholog gene positions within homologous chromosomes reveals that pairs of larger genomes spanning clade boundaries (b-b) show less mixing than smaller genomes (s-s). See also Figures S10 and S11 for pan-metazoan comparisons.

To assess the constraint on the expansion of intergenic distances on linkage at the whole-genome level across multiple species, we first conducted a comparative analysis of genomic distances between orthogroup pairs in 10 representative metazoan genomes (Methods). With 5127 orthogroups shared across these species, we were able to test the correlation between the genome size and pairwise distances in cnidarians, using bilaterians as outgroups. To focus on linkages that are ancestral in all metazoans, we required that at least 5 bilaterian genomes have orthogroup pair distance of maximum 2.5 Mb and we profiled the respective ortholog distance in the cnidarian genomes (1963 such orthogroup pairs were identified). We find that the hydra genome shows the smallest variance of normalized median orthogroup pair distances (**Figure S9**), compared to the smaller genomes of *Hydractinia* (483 Mb) and *Nematostella* (269 Mb). Orthogroup pairs that are linked in bilaterians are thus more likely to stay at a relatively close distance in hydra compared to the other two cnidarians. This trend is retained for orthogroup pairs that are within 1 Mb and 2.5 Mb in bilaterians (**Figure S9**), but no difference was observed for larger and thus likely ancestrally unconstrained orthogroup pairs (e.g., for 10 Mb distance in bilaterians, **Figure S9**). To complement this analysis, we also profiled orthogroup pair genomic distance scaling between *Hydractinia*, *Nematostella*, and hydra. While the genomic distances in hydra are larger and scale with the overall genome size, the relative distance (normalized by the genome size) is smaller in hydra than in *Nematostella*, with the ratio of medians of the relative distances being 0.93 (**Figure 5**c). This suggests that there is generally a higher constraint on local linkages in hydra compared to *Nematostella*.

Beyond pairwise orthogroup distance assessments as a function of the genome size, we sought to assess the degree of positional mixing holistically within chromosomes. For this we have implemented a measure of braid entropy, as initially described in (15). To correct for evolutionary distance biases, we focused on inter-clade comparisons (**Figure 5e**). We find that on homologous chromosomes, mixing entropy is highest for species with smaller (“s”) genomes, in particular between the cnidarian *Nematostella vectensis* to the mollusc *C. squamiferum* (“s-s”, 0.3 Gbp to 0.4 Gbp, **Figure 5**e). Larger (“b”) genomes, such as hydra and the mollusc *M. coruscus* showed less mixing despite the same inter-clade evolutionary divergence (“b-b”, 0.9 Gbp to 1.6 Gbp, **Figure 5**e, **Figure S10**). We have expanded this locus-specific analysis to whole-genome data of over 500 animals and were able to identify similar trends of decreased mixing entropy correlated with the expanded genome size in most of the animal clades (**Figure S11**). Interestingly, the only reversal of the trend, i.e., larger genomes contribute to more mixing, was observed in dipterans. We hypothesize that extensive chromosomal FWM resulting in syntenic dispersal (described as “consolidation” in (75)), together with gene loss, has contributed to this pattern.

Together, both simulation and empirical genomic data suggest a general trend that an increase in genome size correlates with stronger syntenic preservation at the sub-chromosomal, i.e., local or micro-syntenic linkage, level, as well as generally with less mixing on the homologous chromosomes. This finding is mechanistically corroborated by the observed overlap and complex landscape of long-range chromatin loops in hydra bulk and single-cell conformational data.

## Discussion

Our results show that genome expansion is associated with increased length of regulatory contacts and the presence of larger chromatin loops. We provide evidence that this enlargement of DNA conformational contacts, and their evolutionary mixing in particular, is associated with retention of syntenic sub-chromosomal linkages. While not necessarily implying functional co-regulation or co-expression, our findings provide support for the role of multiple enhancer-promoter contact mixing *per se* as a driving force to establish constrained sub-chromosomal evolutionary and regulatory environments (14).

This data provides key insights into the role of regulatory contact mixing in maintaining ancestral genome architecture in animal genomes. It shows how expansion and contraction of a genome forms an orthogonal pillar to other key topological processes, in particular chromosomal fusions (“consolidation”) and inter-chromosomal translocation (“dissociation”) (75). Integrating both pillars, we can project this evolutionary framework onto observed syntenic states in several metazoan clades. For example, we can infer that extensive fusion-with-mixing and the resulting karyotype reduction, which already obscures ancestral chromosomal synteny (“consolidation”, e.g., in some large siphonophore genomes (76)), can be followed by extensive genome contraction and thus more accelerated mixing resulting in syntenic loss, forming a likely mechanism that resulted in some of the most derived genomic architectures such as ascidians (9) or nematodes (77). Genome expansion is, on the other hand, predicted to preserve and decelerate the macro-evolutionary mixing on the newly fused chromosomes. Hence, genome expansion even in the presence of translocation-rich history or whole genome duplications (“dissociation”), helps preserve ancestral gene linkages and regulatory landscapes, as is observed for many and in particular very large vertebrate genomes (6, 10). By careful examination of the observed syntenic patterns, we can thus start dissecting the order of evolutionary events and processes.

Our conformational data on whole-body (Micro-C) and the i-cell lineage single-cell level (Dip-C) resolution gives additional unique insight into both the types and variation of chromatin loop contacts. Single-cell Hi-C data identifies different modalities (principal components) associated with dynamic loop formation within the i-cell stem cell lineage. These results are also validated by the application of DNA FISH for specific genomic loci in native hydra cells. Comparison to bilaterian genomes suggests additional role for CTCF-assisted insulated compartments in these animals that may facilitate emergence of novel regulatory contacts followed by regulatory mixing and entanglement (14) within these regions, introducing additional constraints against dispersion of these loci.

Our study paves the way for further regulatory assessment of the evolutionarily and developmentally dynamic genome of hydra, providing a generalized framework for pan-metazoan comparisons to study association of regulatory entangled states with the origin and evolutionary maintenance of its phenotypic traits, such as stem cell identity and differentiation (78–80).

## Materials and Methods

### Animals culturing

Polyps of Wild-type *Hydra vulgaris* strain AEP and transgenic AEP line expressing cnnos1::eGFP and actin::dsRED (62) were maintained in the animal facility at University of Vienna as described in (81).

### Micro-C library preparation and analysis

The Micro-C library was prepared using the Dovetail Cantata Bio micro-C kit (Cat. no. 017794) following manufacturer protocol. In brief, chromatin was fixed with disuccinimidyl glutarate and formaldehyde, digested with micrococcal nuclease, then extracted by SDS lysis. Fragments were captured using Chromatin Capture Beads, end-repaired, ligated to biotinylated bridge adapter and proximity ligated. After crosslink reversal and protein digestion, DNA was purified and converted into an Illumina-compatible sequencing library using adaptors. Prior to PCR amplification biotin containing fragments were enriched using streptavidin beads. The libraries were sequenced on NovaSeq X (10B XP flowcell, paired end 150 bp).

Micro-C reads were aligned to *Hydra vulgaris* AEP T2T genome (GCF_038396675.1) using Chromap v0.2.7 (82) with `--preset hic` setting, PCR duplicates removal and minimum mapping quality ≥10. Contact matrix was generated with `cooler v0.9.3 cload pairs` (83) at 1kb resolution and then a multi resolution file was generated with `cooler zoomify` with `-- balance` normalisation (83). Additionally, `.hic` file was generated using Juicer tools (84) for loop calling and quick visualisations.

### Micro-C/Dip-C loop analysis

To cover different sizes of loops two loop calling tools were used for Micro-C data. Mustache (44) with settings: `-r 100000, -d 15000000, -pt 0.01` and chromosight (43) with settings: `detect --pattern loops --threads 8 --pearson 0.5 --max-dist 15000000`, at resolution 50000. Afterwards unique pairs were filtered using bedtools pairtopair (85).

For epigenetic data analysis was performed following (36). The raw ATAC-seq and CUT&Tag reads (3 replicates each) from Cazet et al. (36) were mapped to *Hydra vulgaris* AEP genome using chromap (82) with `--preset atac` and `--preset chip` respectively. Alignments were then sorted and indexed with samtools v1.18 (86). For peaks calling in ATAC seq data deepTools v3.5.6 (87) alignmentSieve was used to shifting the reads then MACS3 v3.03 (88) was with following settings: `macs3 callpeak -f BAMPE -g 9.1e8 -n ATAC_1 --outdir peaks --nomodel --keep-dup all -q 0.01`) for calling peaks in all 3 replicates. A consensus peak was defined as the region present in 2 of 3 replicates. Candidate enhancers were defined as H3K4me1+ regions overlapping ATAC-seq consensus peaks but not overlapping H3K4me3 or H3K27me3 peaks. Active promoters were defined as H3K4me3+ ATAC+ peaks within 2kb TSS. All intersections and substractions were done with bedtools. For validation of chromatin states chromatin state segmentation was performed using ChromHMM (v1.25) (89) with 10 state model at 200 bp resolution. Co-occurrency of states was used to validate intersection-based enhancer/promoter calls.

Overlap of each anchor with genomic features (ATAC-seq, H3K4me1, H3K4me3, H3K27me3, active enhancers, and active promoters) was assessed using `bedtools annotate`. Fold enrichment was calculated as the percentage of anchors overlapping a given feature divided by genome wide coverage of that feature.

Motif discovery was performed using HOMER v5.1 findMotifsGenome (90) at anchor ATAC peaks to narrow down range of loop `anchorsRepeat` masking `-mask` and peak coordinates were used (`-size given`). In order to distinguish loop-specific motif enrichment from general properties of open chromatin we compared two genome wide backgrounds with genome wide ATAC-seq peak background. Top-ranked de-novo motif was independently tested by counting in anchor associated peaks versus genome wide peaks using bedtools.

### Micro-C data visualisation

Triangular Micro-C contact matrices were visualised using pyGenomeTracks v3.9 (91). ICE-balanced contact frequencies stored in .mcool format were plotted at 100kb (whole chromosome) and 50kb (regions) resolutions with “Reds” color scale from 0 to 0.01 (normalised contact frequency). Additional annotation tracks were displayed below the matrix using file-types “links” and “bed” of pyGenomeTracks config. This included chromatin loops, loop anchors, ATAC-seq peaks, `ACCCGA` motif, active enhancers and promoters, as well as Wnt genes and oligopool probes locations.

Whole genome and other square matrices were visualised using hicPlotMatrix (HiCExplorere v3.7.6; –log1p –clearMaskedBins –colorMap Reds) (92)

### Dip-C library preparation and analysis

1000 *Hydra vulgaris* AEP animals containing transgenic i-cells with cnnos1::eGFP+actin::dsRED (62) were collected in a 250mL shallow beaker filled with 40mL of *Hydra* medium, and washed to remove food (*Artemia*) and debris. Pronase E was added to the solution at a final concentration of 50 DMC-U ml-1 (93), and rocked in the middle of a linear shaker at a period of 2 Hz at room temperature. When the animals had all detached from the shallow beaker the animals and media were transferred to a 50mL Dounce homogenizer. The animals were disrupted with the tight pestle (B) with 15 complete strokes. The solution was brought to 50mL with additional *Hydra* media, and freshly opened paraformaldehyde was added to a final concentration of 2%. The cells were fixed for 10 minutes at room temperature for 10 minutes, then were quenched with a final solution of 5% BSA on ice for 10 minutes. The cells were filtered with a 35µm filter, centrifuged at 1000 g for 5 minutes at 4 °C. The pellet was resuspended and washed in 5mL of 1% BSA in PBS, then pelleted again. The cells were shipped in 30µL of remaining 1% BSA in PBS to Stanford University on dry ice. The cells were resuspended in 1% BSA in PBS, were separated back into single cells by passing the solution through a 31G needle 10 times, then filtered through a 35µm filter, then counted. One million cells were digested with 80 U of DpnII (`AATT`) given the AT-rich genome of *Hydra*, then ligated (42, 63). The digestion and ligation were confirmed through gel electrophoresis. Digested+ligated nuclei were stained with DAPI, gated for singletons exhibiting RFP-, GFP+, DAPI+, then sorted into 384-well plates at the Stanford Shared FACS facility on a BD S8 (gating figure). Individual cells were prepared in a standard Dip-C library (42, 63) with *Hydra*-specific optimizations of 2.5 µL of Qiagen Protease (QP) per mL of lysis buffer, and 1.25 µL of Tn5 mix (4 mg / mL Tn5 stock) per mL of transposition buffer. Each library was amplified through indexing PCR for 18 cycles using unique dual indices (UDIs) (42, 63). Libraries from one 384-well plated were pooled, size-selected with a 1.20x SPRI cleanup (AMPure XP, Beckman Coulter, A63881), then sequenced on a single Illumina NovaSeq X 10B flow cell on 2 x 150 mode.

Samples were demultiplexed from the raw FASTQ files using fgbio v3.0.0 (94) `DemuxFastq` with reading structure `21B 15B +T +T`, allowing for up to 2 no-calls (`--max-no-calls 2`), 1 mismatch (`--max-mismatches 1`) and mismatch-delta ≥2 (`--min-mismatch-delta 2`) retaining sample-barcodes producing 384 per-cell paired-end FASTQ libraries. After demultiplexing each cell reads were aligned to *Hydra vulgaris* reference genome (NCBI assembly GCF_038396675.1) using BWA-MEM v0.7.19 (95) with `-5SP` flag and then sorted using samtools v1.18 (`sort -n`). Contact pairs were extracted, sorted and deduplicated using pairtools v1.0.3 (`parse –walks-policy 5unique --add-columns mapq, sort, dedup –mark-dups`), retaining only unique read pairs. Pairs file was indexed with pairdix v0.3.9. The valid contact pairs were then converted to Hi-C contact matrices using Juicer Tools (84) v1.22.01 pre.

For analysis single cell Hi-C matrices were normalised using Fan-C v0.9.28 (96)(fanc hic -n), then normalised data from Chromosome 6 (NC_088925.1) was extracted `fanc dump` at default resolution (50kb bins). For each cell contacts were CPM-normalised (count • 10000/total contacts) and after excluding 15 low coverage cells PCA was performed using base R prcomp. Loadings and principal components were visualised using base R and ggplot2.

For visualising 3D chromatin arrangement differences between whole polyp cells and i-cells Micro-C in .mcool format and aggregate of all single Dip-C cells in .mcool format were compared using Fan-C (96) (fanc compare) using -l, normalisation (-Z) and observed/expected approach (-e) and then visualised with pyGenome tracks (91).

### Single cell loop calling

For each cell ‘`.pairs`’ from single cell Dip-C data were converted to SnapHiC contacts format (`chr1/pos1/chr2/pos2`), with one contact per line. Loop calling was performed in 3 steps using SnapHiC pipeline (97) with bin size being 10kb for all steps; Firstly, we did imputation with the inverse RWR method (`--steps bin rwr`), 7Mb distance (`--dist`). Then interaction calling (`--steps interaction`) and post processing (`--steps postprocess`) were performed with 6Mb distance yielding per-chromosome `.bedpe` files (loop anchors and summit coordinates). These per chromosme files were filtered using `z > 1.96` threshold to derive per-cell per-chromosome loop `.bedpe` files. Then using bedtools and base python statistics and plots were generated.

*Hydra viridissima* Hi-C data visualisation

Publically available Hi-C data for *Hydra viridissima* has been acquired using accession ERX12760466, it was mapped to jhHydViri1.1 (GCF_964215635.1) reference genome in the same way as described for *H.vulgaris* Micro-C data.

### Enhancer-promoter contact prediction

To identify the candidate genomic regions that are regulating the genes in the *Hydra vulgaris* AEP T2T genome assembly, we used Activity by Contact model (47) following the pipeline described in (98). Data used in the analysis included the Micro-C-seq, ATAC-seq, and H3K4me1 CHIP-seq data described in *Micro-C/Dip-C loop analysis* section in the methods, and RNA-seq (Acession.No:SRX14496578) all generated from the AEP strain. RNA-seq data was downloaded using SRA-Toolkit (99) from NCBI Sequence Read Archive (100). Reads were mapped to the reference genome using STAR version 2.7.11b (101), and the gene counts were calculated using Rsubread (102). ATAC-seq peaks were called using MACS3 version 3.0.4 (88) and sorted using BEDtools version 2.31.1 (85). Hi-C interactions were extracted from .hic into .bedpe format using hictk version 2.2.0 (103). SAMtools (86) was used to rename the chromosomes in `.bam` files when needed to match the naming to the NCBI release, and to index the `.bam` files. Enhancer predictions with an ABC score above 0.05 were used. ABC score is a normalized estimate of how much a candidate enhancer contributes to a gene’s regulation, based on enhancer activity and enhancer–promoter contact, and values of minimum 0.015 or for higher confidence 0.02 have been suggested as significance thresholds (104).

### Conserved region analysis

Whole genome alignments for whole genome comparisons of 9 different hydrozoan species (including two different strains of *Hydra vulgaris*, 105 and AEP) with two outgroup species (*Nematostella vectensis*, *Branchiostoma floridae*) were constructed using Cactus (v3.0.1) (105). Used species with genome-assembly accession numbers of the total of twelve species were the following. *Hydra vulgaris* (AEP): GCF_038396675.1, *Hydra vulgaris* (105): GCF_037890685.1, *Hydra oligactis*: GCA_024195425.1, *Hydractinia symbiolongicarpus*: GCF_029227915.1, *Physalia physalis*: GCA_041430235.2, *Hydra viridissima*: GCA_964215635.1, *Ectopleura larynx*: GCA_978473255.1, *Candelabrum cocksii*: GCA_963930725.1, *Turritopsis rubra*: GCA_039566895.2, *Clytia hemisphaerica*: GCF_902728285.1, *Nematostella vectensis*: GCA_932526225.2, *Branchiostoma floridae*: GCF_000003815.2. Since no prior evaluation of phylogenetic relationships or estimations of divergence times between the aligned genome assemblies were done, the (unrooted) guide tree used for the alignment was based on already published phylogenies inferred on mitochondrial *rrnS* (16S rRNA) and *cox1* (cytochrome oxidase I) (106) as well as entire mitochondrial genomes (107, 108). Due to incoherent findings regarding the internal phylogeny of five species of Anthothecata (*H. symbiolongicarpus, T. rubra, C. cocksii*), Leptothecata (*C. hemisphaerica*) and Siphonophorata (*P. physalis*) to a consistently reported monophyletic group of aplanulates (*H. vulgaris, H. oligactis, H. viridissima, E. larynx*) within hydrozoans, their relative placement in the guide tree was simplified into two neighbouring, binary resolved sub-branches. Resulting in a subtree with *H. vulgaris, H. oligactis, H. viridissima* and *E. larynx* adjacent to a subtree with *H. symbiolongicarpus, T. rubra, C. cocksii*, *C. hemisphaerica* and *P. physalis* (with the here listed order from late to earliest splitting species within the subtrees). The subtrees were contrasted with *N. vectensis* and *B. floridae* as outgroups. All internal branch lengths of the guide tree were set constantly to one by Cactus since none were additionally provided as input. The genomes of *Hydra vulgaris* (AEP), *Hydractinia symbiolongicarpus* and *Branchiostoma floridae* were set as Cactus-internal reference genomes to serve as outgroups for the construction of sub-alignments.

With *Hydra vulgaris* AEP strain as the reference genome, resulting multiple alignments were extracted from the whole-genome hierarchical alignment using `cactus-hal2maf` (109). For each Hydra reference chromosome, a neutral phylogenetic model was fitted with PHAST (110) phyloFit under the REV substitution model using the HAL-derived species topology, and sitewise conservation scores were computed with (LRT mode) (111) in Hydra reference coordinates. Per-base phyloP WIG files were thresholded at phyloP ≥ 2, and adjacent high-scoring bases were merged with BEDTools to define candidate conserved regions. PhyloP is a per-base evolutionary conservation score, and a cutoff of ≥ 2 is reasonable because positive values indicate conservation, and a score of 2 corresponds to about p value = 0.01 against neutral evolution. `bigWigAverageOverBed` from Kent tools (112) was used to calculate the average and maximum phyloP scores over sets of regions.

### OligoFISH probe preparation and protocol

#### Primary probes locus specific

Probes targeting loop anchors were designed using OligoMiner pipeline (113) with blockParse l 23 -L 60 -t 32 -T 47 -S 10, bowtie2 stq --no-hd -k 2 --local -D 20 -R 3 -N 1 -L 20 -i C,4 --score-min G,1,4 -S. outputClean.py -T 37 -p 0.999. kmerFilter -m 18 -k 30. structureCheck -T 37 -t 0.1 -s 330. Probes were generated against Chromosome 6 of *Hydra vulgaris* AEP strain (NC_088925.1, assembly GCF_038396675.1). The region containing potential loop anchors as identified by Micro-C was divided into 10kb windows, then for each window OligoMiner pipeline was applied and the window yielding the highest number of oligos was selected for probe synthesis (148 oligos pool for region NC_088925.1:7000000-7010000 and 190 oligos pool for region NC_088925.1:11650000-11660000). Each individual oligo was structured as follows: *5’-[nullomer]-[TT_bridge]-[genomic_sequence]-[TT_bridge]-[nullomer]-3’*. Probes were sequenced at 10 pmol per oligo by Integrated DNA Technologies and resuspended in 25 µL TE buffer (10 mM Tris, 1 mM EDTA, pH 8.0).

### Primary DNA FISH probes for repetitive regions

Centromeric sequences defined in (11) were mapped to the *Hydra* T2T genome assembly using BLAST to identify corresponding centromeric regions. In order to find candidate probes 25-mers were counted in both the centromere candidate sequences and the reference genome using Jellyfish v2.3.1 (114) (-m 25 -C, canonical mode). Genome wide k-mers occurring less than 10 times were discarded (-L 10). Then k-mers were filtered to have at least 100 copies in the genome and the dust score ≤5. Dust score was calculated as sum(ct(ct-1)/2. This resulted in 657 k-mers which were then mapped to the reference genome and windows with the densest repeat signature were used. The genome was then split in centromere and non-centromere regions. Those k-mers that were present outside of centromeres in more than 10 copies were removed from the inside k-mer set using comm −23. Remaining inside unique k-mers were filtered by copy number greater than 100 and dust score ≤15 yielding 1308 centromere specific k-mers. Candidate probes were sorted by genome copy number and dust score. One of the top candidate probes (25-mer CAAAAAAACAGGTTTTCACTGATTT) was present at 7693 genomic copies (98.4% in centromeres) and had a dust score of 13. For designing the probe we added reverse complement of secondary (CACACGCTCTTCCGTTCTATGCGACGTCGGTG) resulting in the following probe: 5’CAAAAAAACAGGTTTTCACTGATTT-CACCCACCGACGTCGCATAGAACGGAAGAGCGTGTG-3 ’.

Telomeric probes were designed to contain 4 times C-rich telomeric repeat CCCTAA, as well as two nullomeres (CGCAGCTCCACTTGATCTCGCTGGATCGTTCT) at both ends separated by CACC bridge: 5’-CGCAGCTCCACTTGATCTCGCTGGATCGTTCT-CACC-CCCTAACCCTAACCCTAACCCTAA-CACC-CGCAGCTCCACTTGATCTCGCTGGATCGTTCT-3’).

Telomeric and Centromeric probes were synthesised at 1.2 nmol and 3.8 nmol respectively by Sigma-Aldrich, upon arrival they were diluted to 100 µM in TE buffer (10 mM Tris-HCl pH 8.0, 1mM EDTA) and stored at −20 °C.

#### Secondary probes

Two types of secondary probes with homology to the nullomer of respective primary probes were designed as described in (113, 115). The secondary probe against oligopool targeting loop anchor around 7Mb locus was designed with two ATTO565 fluorophores at both ends (`5ATTO565N/CACACGCTCTTCCGTTCTATGCGACGTCGGTGtttttttt/3ATTO565N`) and the secondary probe against oligopool targeting locus around 11.6 Mb was designed to carry FAM at 5’ end (5FAM/AGAACGATCCAGCGAGATCAAGTGGAGCTGCG). They were synthesized by Microsynth (60.5 nmol) and Gene Script (5 nmol) respectively and resuspended in TE buffer (10 mM Tris-HCl pH 8.0, 1mM EDTA) to obtain a 100 µM that was aliquoted and stored at −20 °C in the dark until use.

### DNA FISH Sample preparation and Staining

Samples were prepared following modified protocol from (116). *H.vulgaris* AEP strain were starved for 2 days, collected in 1.5 mL tube and fixed with 4% Paraformaldehyde in PBS for 10 min. After fixation animals were washed three times with 1X PBS and 2-3 polyps were placed on poly-L-lysine coated slides (cat. No. D8920) in a PBS drop. To obtain a near single layer of cells, coverslips were placed on the sample with constant pressure then removed by brief immersion in liquid nitrogen and lifting with a scalpel.

Staining was performed following a modified protocol from (117). After the samples returned to room temperature they were washed twice in 1X PBS for 5 min, then permeabilised with 0.5% Triton X-100 (cat. No. 93443) PBS for 10 min and washed twice with PBS. Slides were then incubated in 0.1 M HCl for 10 min, followed by incubation in freshly prepared 0.0025% pepsin (Sigma-Aldrich, cat. No. P6887-1G) in 0.01 M HCl solution at 37°C for exactly 2 min. Subsequently, samples were washed three times in 2X SSCT for 1 min, then in 2X SSCT+50% formamide for 20 min at room temperature and 20 min at 60°C.

Hybridisation was performed in freshly prepared 50% formamide, 2X SSC, 10% dextran sulfate (cat. No. D8906) and 0.5 mg/mL salmon sperm DNA (cat. No. 15632011). Primary probe pools targeting each anchor region (148 and 190 oligos) were added at ∼1.2 and ∼1.5µM respectively (8 nM per oligo), secondary probe pools were added at 2 µM and RNase A was added at 400µg/mL. Hybridisation mixture (∼12 µL) was applied to each sample and sealed with Fixogum (At. no. 29010010000), and denatured at 80°C for 5 min on an aluminum block in a water bath. Slides were then incubated in a humidified chamber at 35°C for 12-18h. Following hybridisation, slides were washed for 15 min in 2X SSCT at 60°C, then for 10 min in 2X SSCT at room temperature and for 10 min in 0.2X SSC. Samples were rinsed in 1X PBS, counterstained with DAPI (1:5000) for 15 min, rinsed again in 1X PBS and mounted in DAPI-free Vectorshield (cat. No. H-1000-10).

Confocal imaging was performed at the core facility of Cell Imaging and Ultrastructure Research (CIUS) at University of Vienna using Leica Stellaris 5 confocal microscope equipped with HC PL APO 63x/1.30 GLYC objective and HyD S detectors. Three channels were collected sequentially with a pinhole set to 1 AU and a z-step of 0.15-0.5 µm. FAM was excited using OPSL 488 laser at 15-30% power (detector gain 55-107) and ATTO565 using OBIS 561 at 8.5% power and detector gain 15.1. DAPI was excited using a 405 nm diode laser.

### Telomeric and centromeric FISH

Telomeric and centromeric DNA FISH was performed following the protocol described above. Primary probes were used at 2 µM for telomeric and centromeric probes each. Secondary probes were used at 2 µM each as well. Microscopy was performed as described above with a z-step of 0.3 µm, OPSL 488 at 10-20% power, detector gain 9.3-52.6 and OBIS 561 laser with 3-8.3% power (gain 4.1-8.9).

### Orthology construction

Orthologous groups (OGs) were identified using OrthoFinder v2.5.5 (118) with default parameters. Raw GFF from NCBI were cleaned up to contain only the longest isoforms by using AGAT v1.6.1 (119) agat_sp_keep_longest_isoform.pl. Then gene coordinates were extracted from GFF3 flies of respective genome assemblies and linked to OGs. Firstly, orthologs were filtered to retain only those that are present in more than 5 bilaterian and at least 1 cnidarian species. Then orthologs were parsed to create pairs of orthologs (OG pairs). OG pairs were retained if their maximum distance in any of the seven bilaterian species did not exceed a specified threshold (e.g., 2.5Mb on the same chromosome). Then the table of OG pairs was updated with chromosomal scaffolds for each species having respective OG pairs. Statistical analysis and visualisation were performed using base R v4.5.2 and Python v3.9.21.

### Hox genes synteny analysis

Protein sequences of Hox genes for selected species (*B.floridae*, N.vectensis, H.vulgaris, H.symbiolongicarpus, M.musculus) were obtained from NCBI. An initial dataset of 107 sequences was aligned with MAFFT v7.526 (120) `–auto –anysymbol` and phylogeny was inferred using IQ-TREE v2.4.0 (121) `iqtree2 -B 1000`. The resulting tree was used to identify and remove non-Hox sequences and to collapse same locus sequences, duplicates were further cleaned up by using `tblastn` from BLAST+ v2.16.0 (122) resulting in a dataset of 81 sequences. This filtered set was re-aligned with MAFFT and phylogeny recomputed under the same settings. After that Hox genes were grouped and their genomic position was found using `tblastn` against respective chromosome-level assembly. Best scoring hits per query were kept and sequences mapping to the same genomic location were collapsed to a single representative. Synteny was further visualised using pyGenomeViz v1.6.1 (123).

### Wnt gene annotation and assignment

Wnt protein sequences were obtained from NCBI. Sequences were aligned with MAFFT v7.526 (120) and phylogeny was inferred with IQ-TREE v2.4.0 (121) to assign each Wnt sequence to the gene family. Genomic positions were determined by using best hit of BLAST+ v2.16.0 (122) `tblastn` against respective chromosomal level assembly. Synteny was visualised with pyGenomeViz v1.6.1 (123).

### Braiding analysis

For the calculation of entropy across multiple species, *Branchiostoma floridae* (amphioxus) (GCA_000003815.2) peptides were mapped using miniprot (v0.12) to multiple Genbank assemblies. The resulting files were converted into .chrom format for easier handling. For pairwise comparison, .chrom files were merged into combined psynt files using a Python script (Python 3.13). Using R (v4.5.1), pairwise gene locations and a matrix of chromosomal homology significance were generated to identify significant combinations. For entropy calculation in MATLAB (R2025b), the Braidlab package (124) was used. For each significant syntenic block, the relative order of homologous genes along the chromosomes of species A and species B was extracted and treated as two trajectories in gene-order space. These trajectories were converted into braid objects and braid entropy was calculated to quantify gene-order differences between species; low entropy indicates conserved gene order, whereas higher entropy indicates increased genomic rearrangement. For pan-metazoan braiding analysis, amphioxus peptides were mapped to 607 species (**Table S1**). Braiding was then computed between *Nematostella*, *Hydra* and amphioxus to their respective outgroup species with the braidlab package as described above. Results of braid entropy were plotted with R.

## Data, Materials, and Software Availability

All data is deposited at PRJNA1447980. Enhancer-promoter contact predictions, and the conserved regions across 10 species in the Hydra vulgaris genome AEP strain T2T assembly are available at: 10.5281/zenodo.19388751.

## Supporting information

Supplementary Table S1. List of species used in mixing (braiding entropy) analyses. NCBI tax id is included in the species name.

## Acknowledgments

We thank Charles David for giving feedback on the manuscript and hydra confocal imaging and Arnau Sebe-Pedros for comments and feedback on the manuscript. We thank all the members of Simakov group for support and discussions through this project and Mirela Król for assistance with figures. We are thankful to the member of Department of Neuroscience and Developmental Biology (University of Vienna) for great and collaborative environment, among them Sanjay Narayanaswamy, Juan Daniel Montenegro Cabrera, Grigory Genikhovich, Daria Filimonova for their experimental and computational suggestions, as well as to Angela Caballero Alfonso and Samruddhi Kankekar for assisting in animal maintenance. We thank Thomas Steinacker and Brian J. Beliveau for giving suggestions on computational and experimental aspects of DNA FISH. Y.T., T.R., D.T.S., F.S., O.S. were supported by the European Research Council’s Horizon 2020: European Union Research and Innovation Programme, grant no. 945026. L.T. was supported by BWF CASI, Sanofi, Baxter, PSF MIND, McKnight, Rita Allen, Wu Tsai/Knight, Bio-X, and RCSA. S.T. and B.P. were supported by Barres. F.B. was supported by CNRS. Computational results of this work have been achieved using the Life Science Compute Cluster (LiSC) of the University of Vienna. Microscopy was performed at the Core Facility Cell Imaging and Ultrastructure Research, University of Vienna - member of the Vienna Life-Science Instruments (VLSI). We thank Brigitte Schmidt and Katy Schmidt for assistance and valuable input on optimising our confocal microscopy.

## Author contributions

Y.T. and O.S. designed the overall study. Y.T. and T. R. performed micro-C. D.T.S., S.T., B.P., L.T. performed Dip-C. Y.T. conducted Micro-C and Dip-C analyses. Y.T. designed and optimised oligopaint probes; Y.T. and F.B. designed the DNA FISH experiments; Y.H, E.A.I. and L.W. performed the experiment. K.K., T.K. identified centromeres. F.Sa. conducted ABC and phyloP analyses, F.St. conducted genome alignments. Orthology and analysis was done by Y.T. Braiding analysis was done by J.G., Y.T., and O.S. Y.T. and O.S. wrote the paper with input from all authors.

## Figures

**Figure S1.**
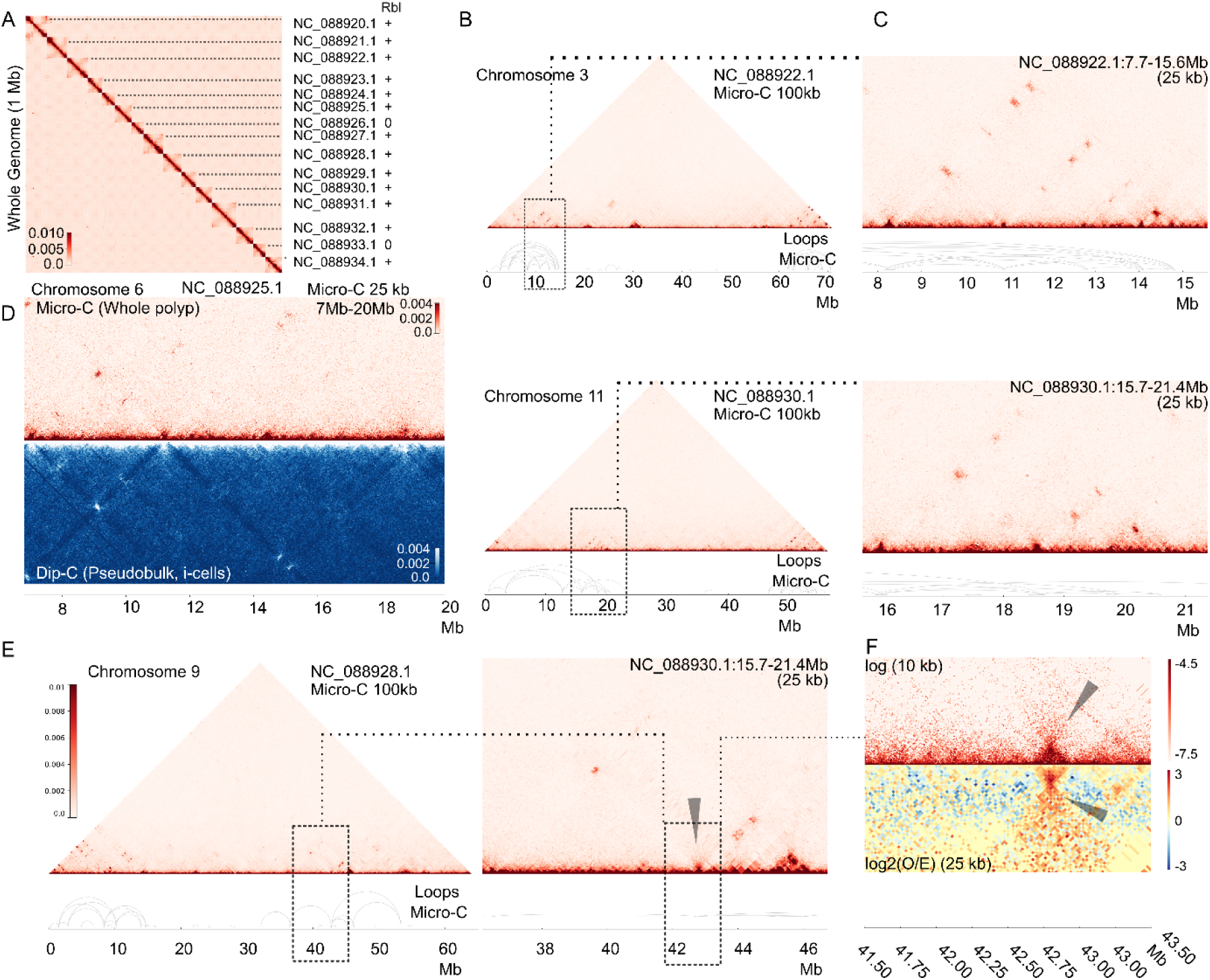
Micro-C data reveals long-range chromosomal interactions in hydra. (A) Whole genome Micro-C data at 1 Mb resolution. Scaffolds are indicated as well as presence of Rabl-lIke configuration is noted in the Rbl column. (B) Micro-C maps of selected chromosomes at 100kb resolution, with higher order hub-like chromatin features highlighted, and predicted loops displayed as separate tracks. (C) Zoom-ins of features of interest on respective chromosomes. (D) Split view of whole genome Micro-C (upper half), Dip-C pseudobulk (merged Dip-C data from all individual cells, bottom half) data at 500kb resolution. (E-F) Example of a single loop of several megabases with signatures of symmetrical jet, suggesting SMC mediated loop extrusion. Zoom-in (E) and observed/expected at high resolution (F, “O/E”, 10kb and 25kb) show jet-like patterns.

**Figure S2.**
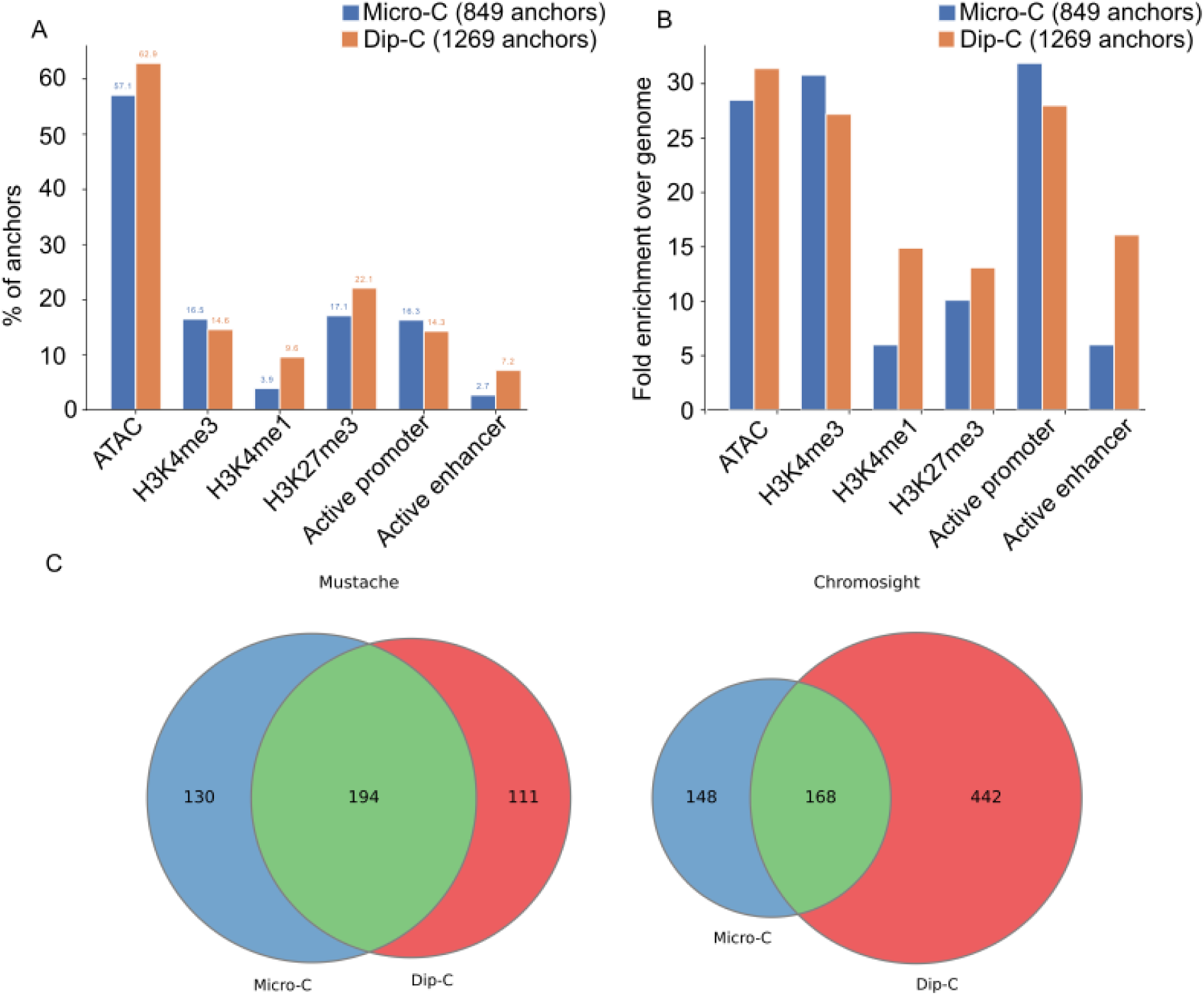
Loop calling comparison and epigenetic marks. (A) Mustache and Chromsight were used to predict chromatin loops in both micro-C and pseudo-bulk Dip-C data. Mustache was more consistent between the two experimental HiC datasets. As both Mustache and Chromsight loops could be validated in the Hi-C maps, we merged the two predictions. (B) Epigenetic chromatin marks in loop anchors in both micro-C and pseudo-bulk Dip-C data (left) and (right) enrichment of marks in loop anchors relative to all other genomic regions.

**Figure S3.**
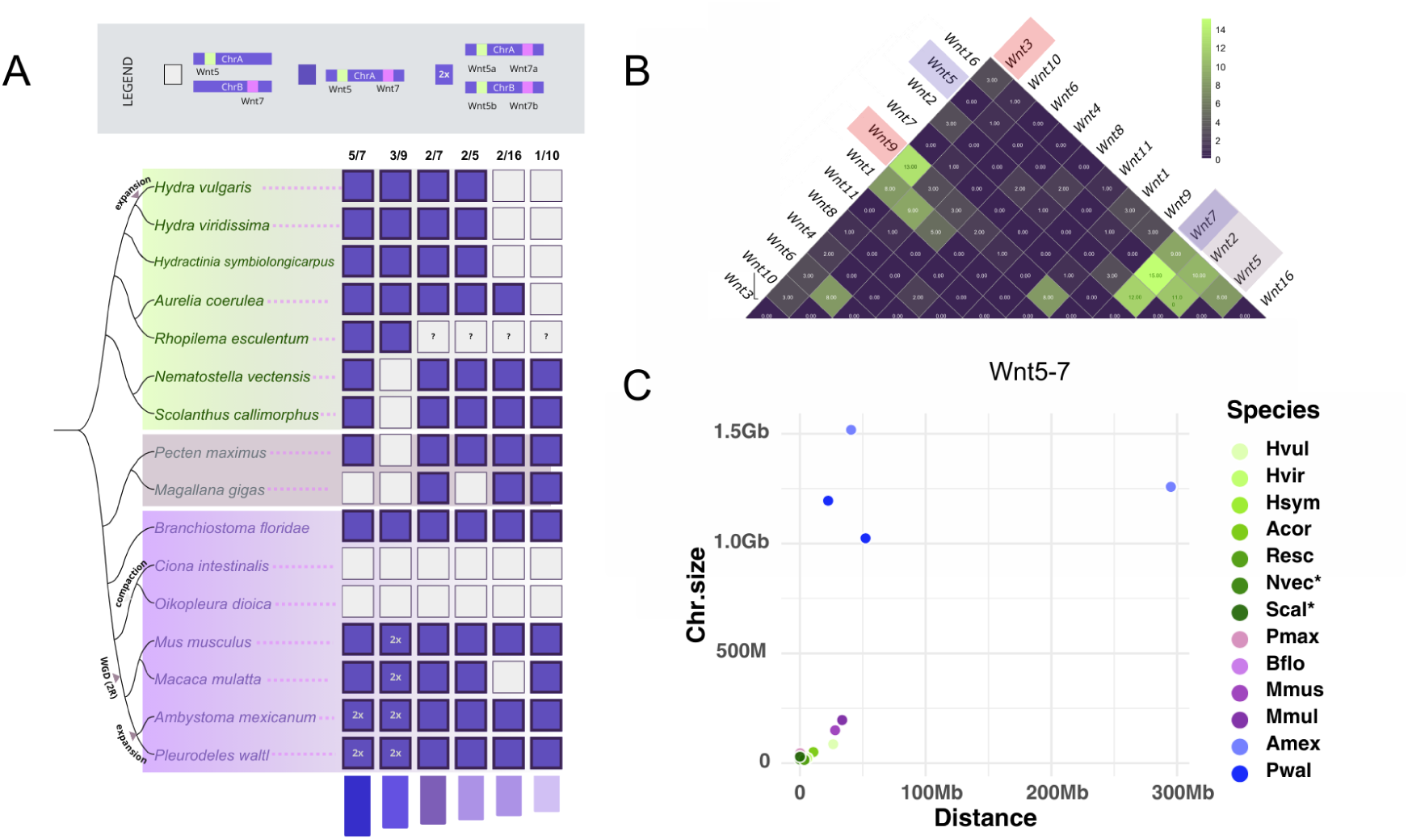
Wnt complement and clustering across selected metazoan genomes. (A) Phylogenetic distribution of Wnt ligand clustering across several representative metazoan species. (B) Average distance between Wnt ligands across species. (C) Size of Wnt clusters in relation to chromosome size for the Wnt5-7 association.

**Figure S4.**
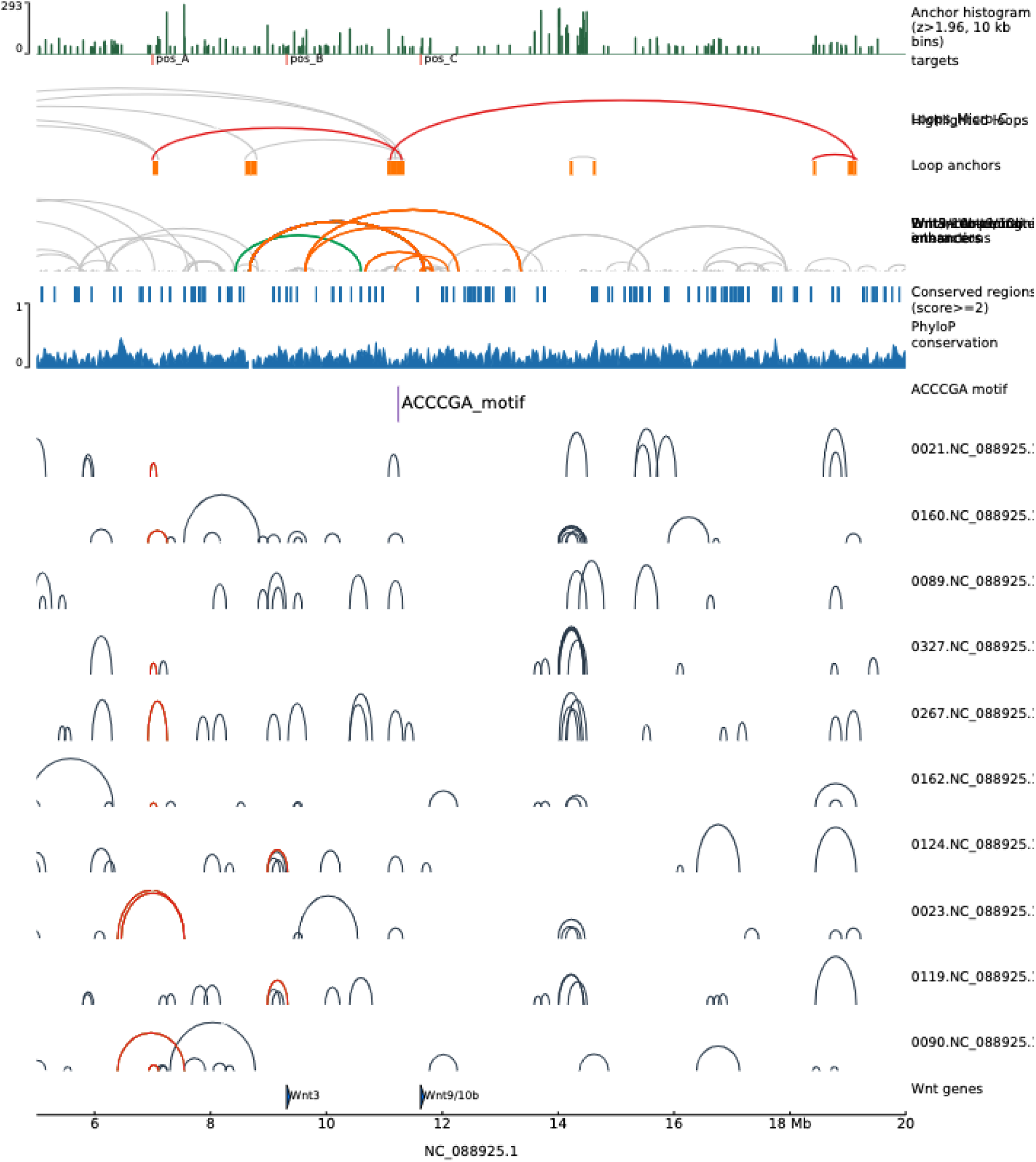
Wnt locus in the hydra genome and its associated chromatin loops and predicted enhancer-promoter contacts. Different tracks represent, from top to bottom: anchor density along the chromosome, predicted loops in Micro-C data, predicted enhancer-promoter contacts, conserved regions, predicted loop anchors, enriched motif location, single-cell Dip-C data loop predictions (for 10 cells, labelled with number corresponding to the unique cell), location of the Wnt genes.

**Figure S5.**
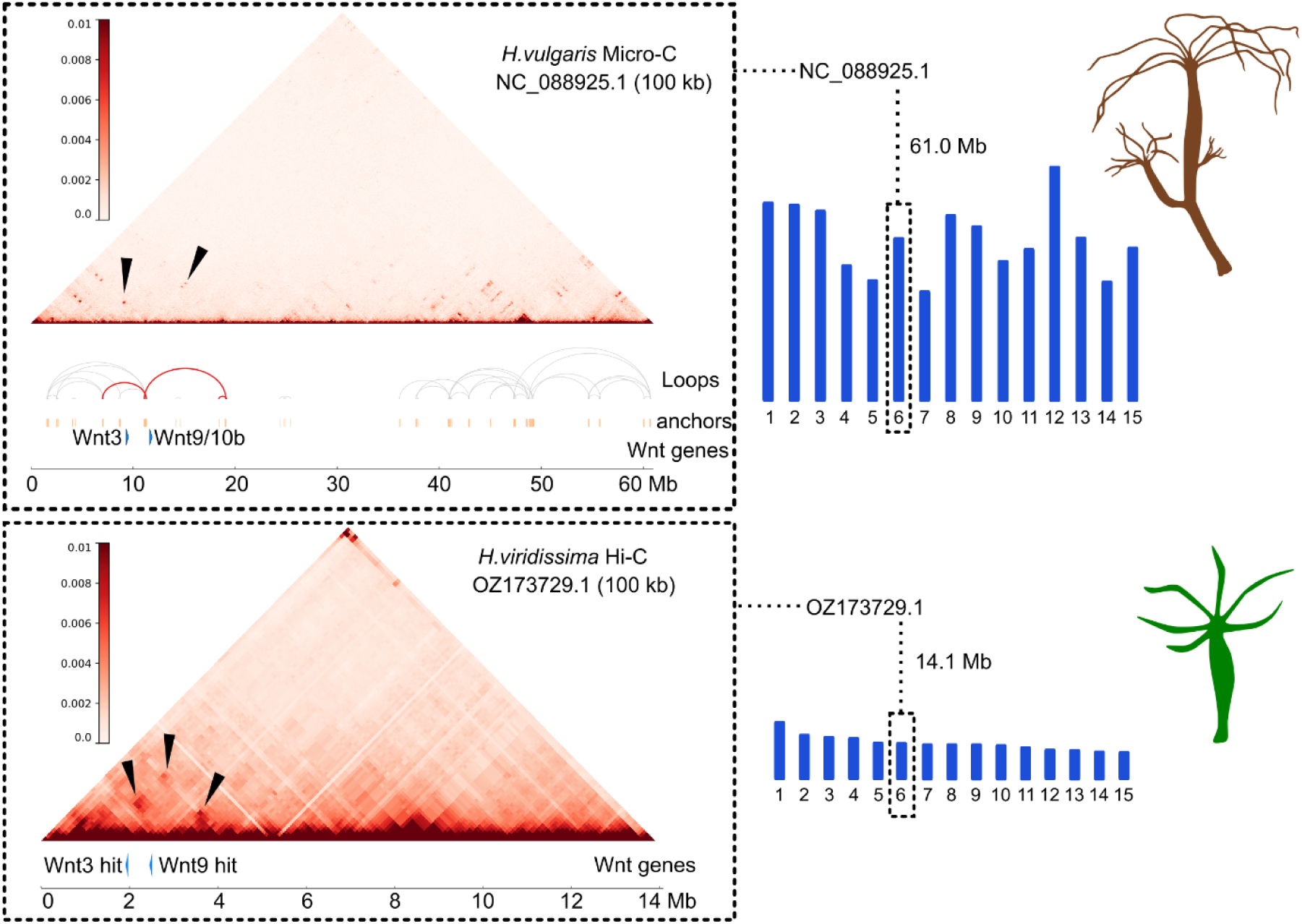
Comparison of the Wnt 3-9/10 locus between *H. vulgaris* and *H. viridissima*. Green hydra chromosome OZ173729.1 (mirrored) with top tBLASTn hits for Hydra Wnt3 and Wnt9/10a,b.

**Figure S6:**
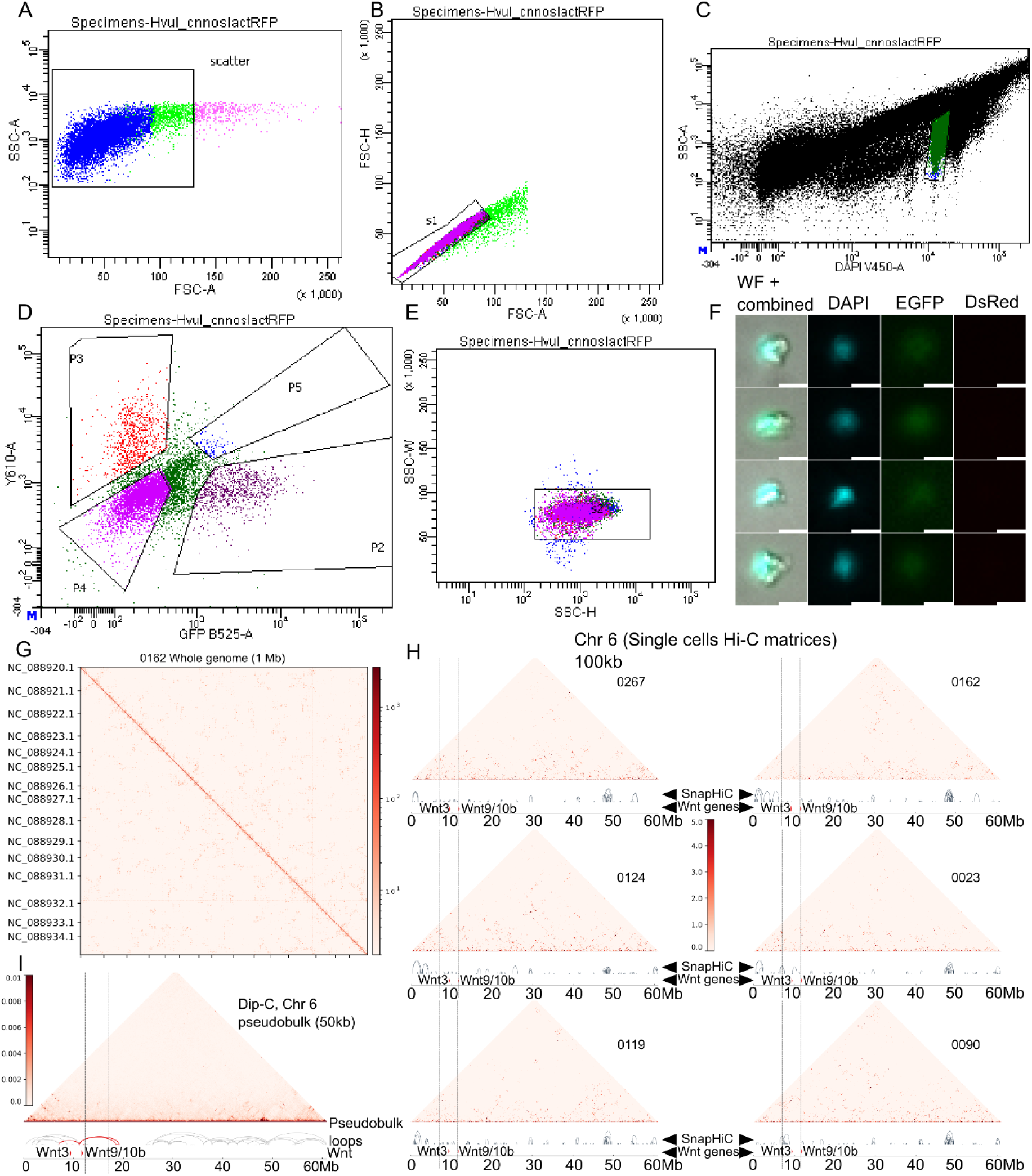
Single-cell gating strategy for Hydra i-cells. Cells were gated with the following strategy (A and B) Clumps were removed with FSC and SSC singlet gates (C, s2) DAPI-positive cells were sorted, and (E, P2) GFP-positive, RFP-negative cells were sorted into a FACS tube (P2 gate), and the cell phenotype of GFP+ and RFP- fluorescence was confirmed on a fluorescence microscope (F). This population of cells was sorted into single wells on a BD S8 FACS machine using DAPI singlet (linear scale) gates. Scale is 10 µm (G) Whole genome Hi-C matrix of a single cell at 1Mb resolution. (H) Additional examples of Chr 6 Hi-C matrices of different single cells at 100kb resolution. (I) Pseudo-bulk of Chromosome 6 at 50kb resolution. For its generation pairs of all cells were combined and mapped to the reference genome. The data illustrates the presence of a loop around Wnt3/9 locus as in Micro-C.

**Figure S7.**
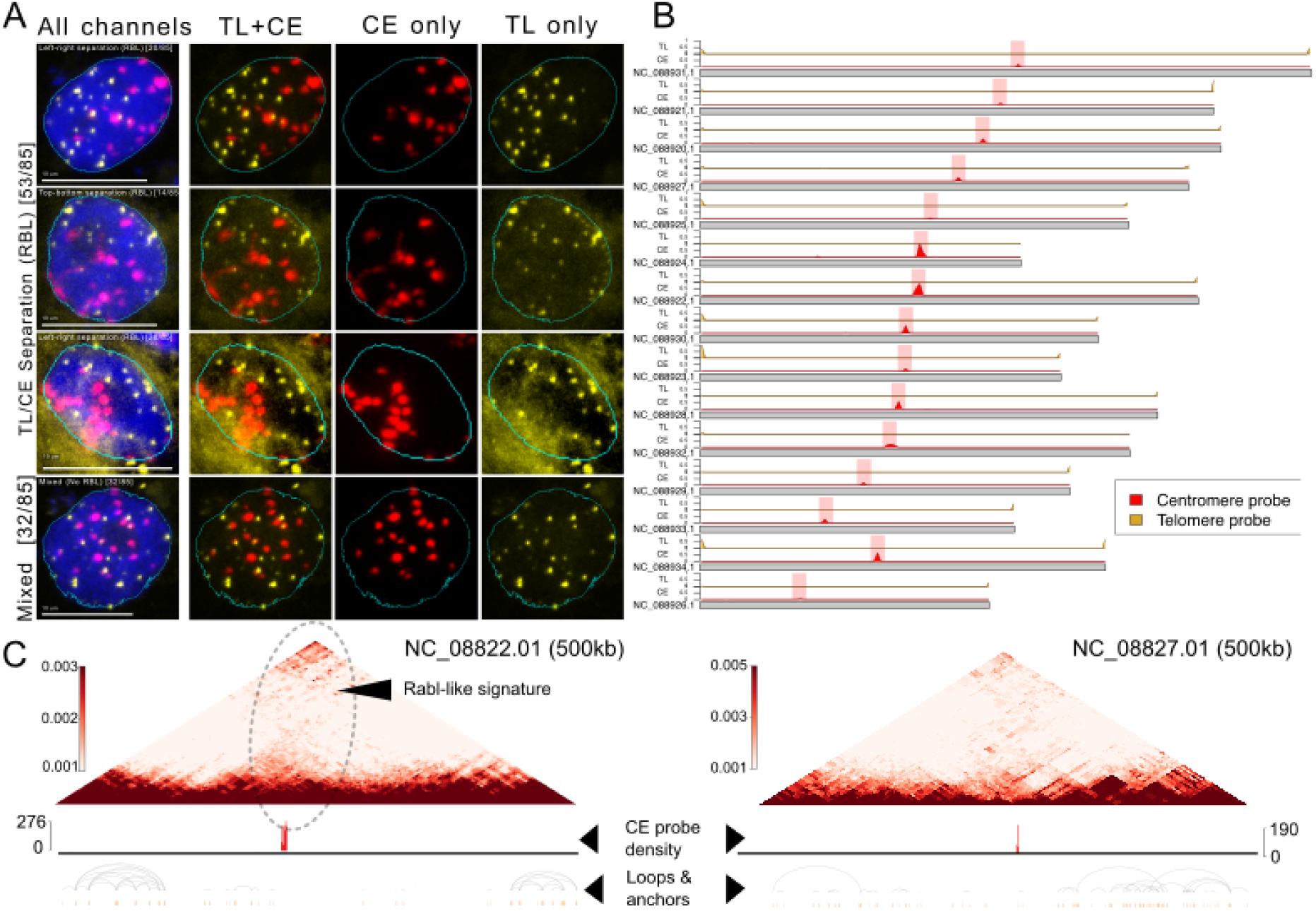
Rabl-like configuration of hydra chromosomes confirmed with DNA FISH and quantification. (A) Staining with CE (centromere) and TE (telomeric) probes. (B) Distribution (exact matches density) of telomeric and centromeric probes along hydra chromosomes. (C) Micro-C maps highlighting the Rabl-like pattern.

**Figure S8.**
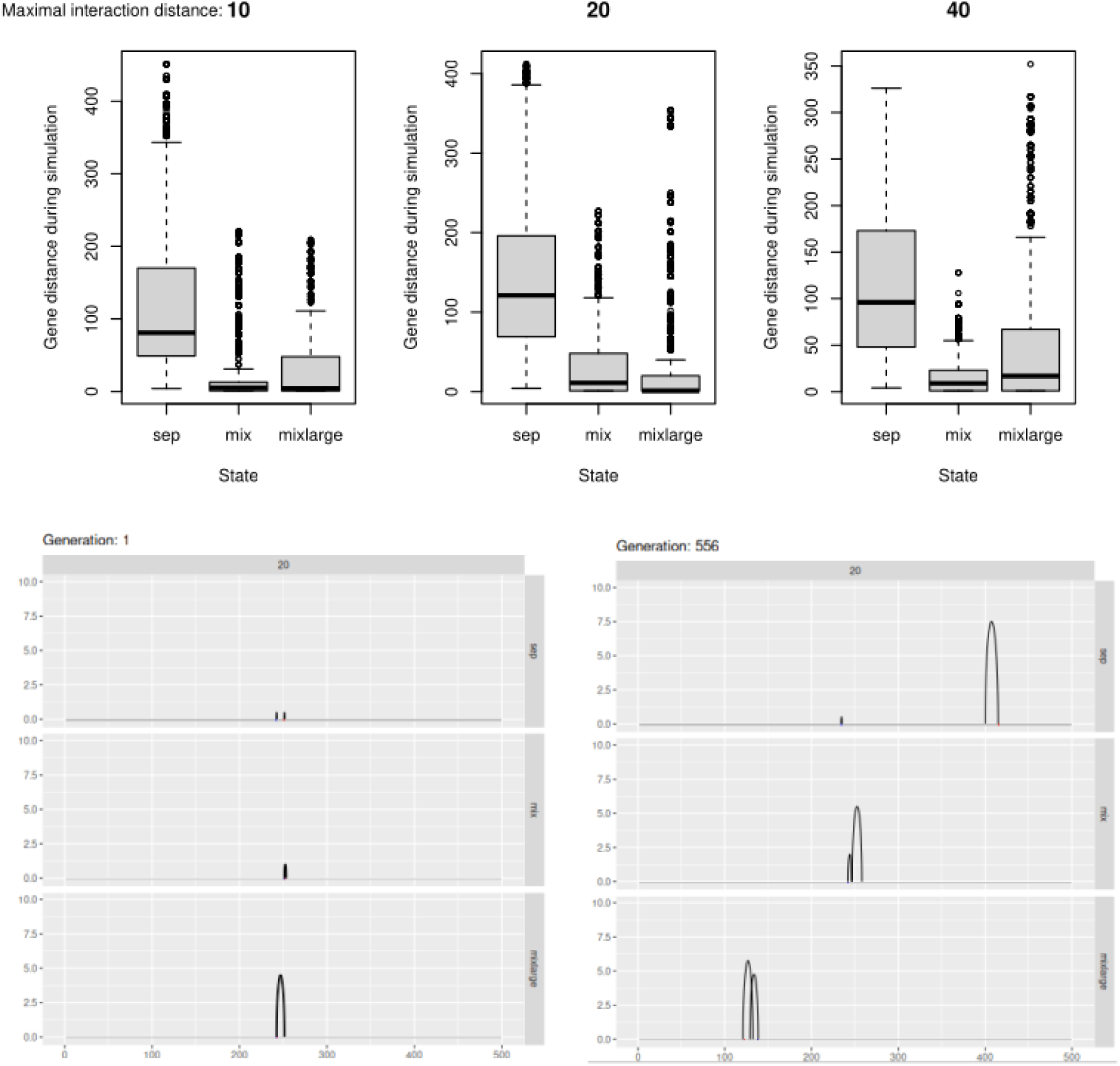
Simulated mixed states are more constrained in expanded genomes. Chromosomes of 500 positions with two enhancer-promoter contacts shown as arches in two possible states (either separated “sep”, mixed “mix”, or mixed and enlarged “mixlarge”) are passed through random inversions (exponential distribution). Only inversions that do not separate individual enhancer-promoter contact beyond the maximal interaction distance of 10, 20, or 40 are recorded and included in the simulation result. In total there were 1000 cycles (generations) per simulation, 5 replicates. Upper row: cumulative summary for the three maximal interaction distances and three possible initial states are shown for the gene distance between the two genes during the whole simulation window. The two genes are substantially constrained in their separation in both mixed and mixed-and-enlarged states. Lower row: Example for one of the replicates at the initial state of the three states and at a later state of the simulation highlighting still neighboring entangled regulatory states.

**Figure S9.**
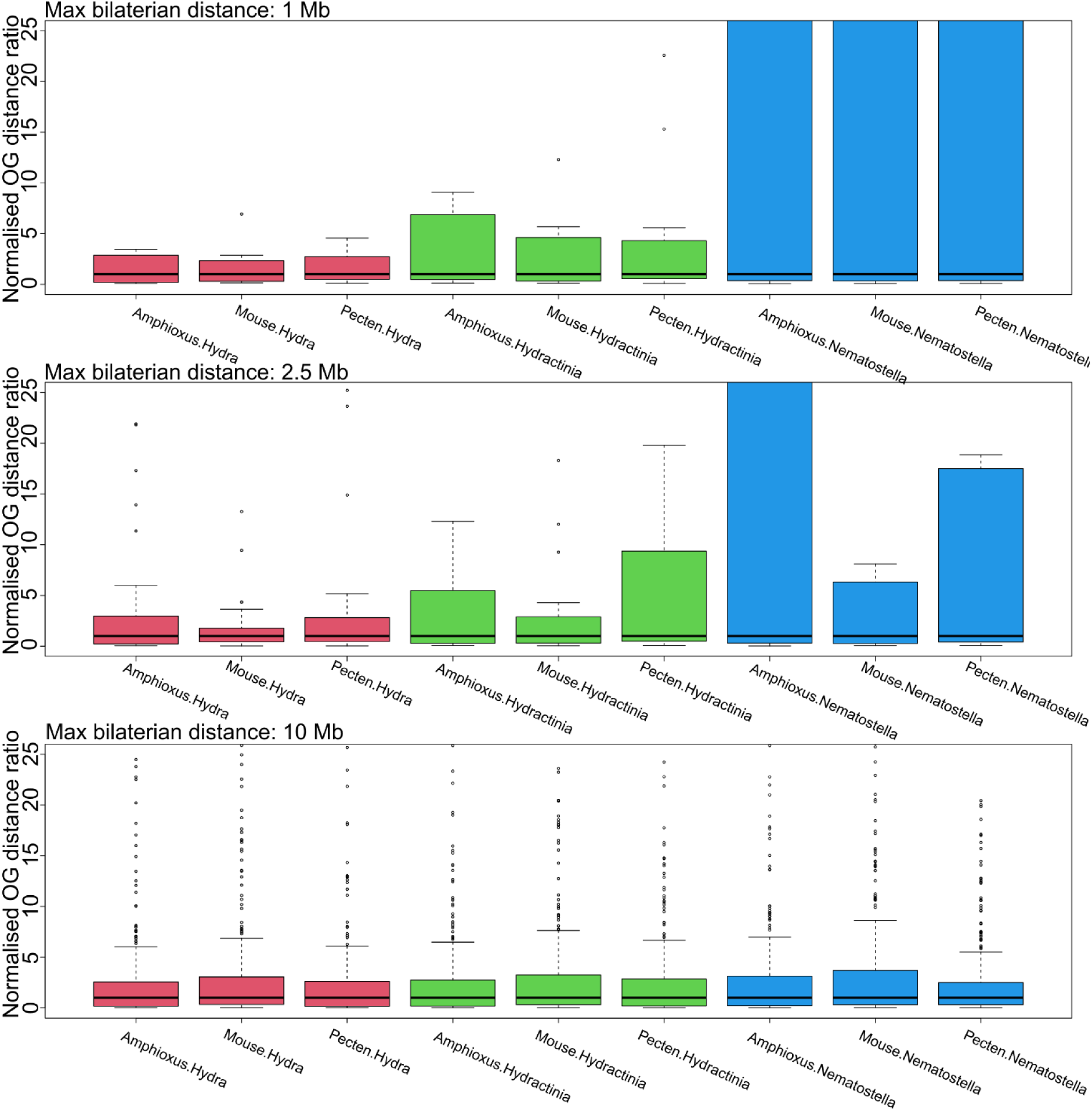
Normalized orthogroup pair distances show less variance in larger cnidarian genomes. The ratio in orthologous gene-pair distances was computed between pairs of species and is normalized by its median to control for the genome size effects. Orthologous gene pairs that are maximally within 1Mb (top), 2.5Mb (middle), and 10Mb (bottom) range in any of the seven profiled bilaterian genomes are used to compute their scaling in Hydra (red), Hydractinia (green), and Nematostella (blue). Only for unconstrained orthogroups (maximal distance 10Mb in bilaterians) no difference in variance is observed.

**Figure S10.**
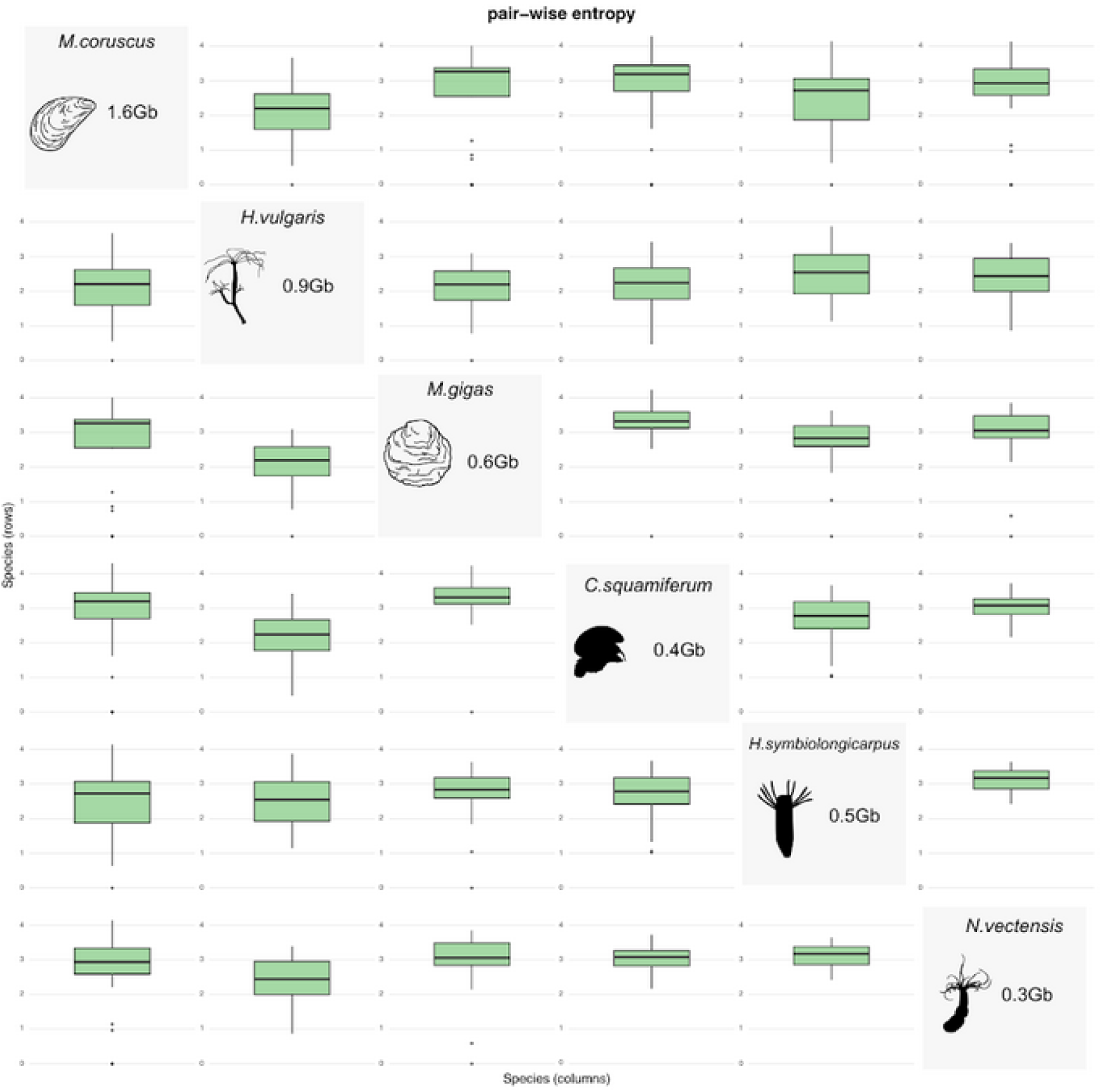
Comparative braid entropy in a select set of species as a function of genome size. Larger genomes show less mixing of their orthologs than smaller genomes.

**Figure S11.**
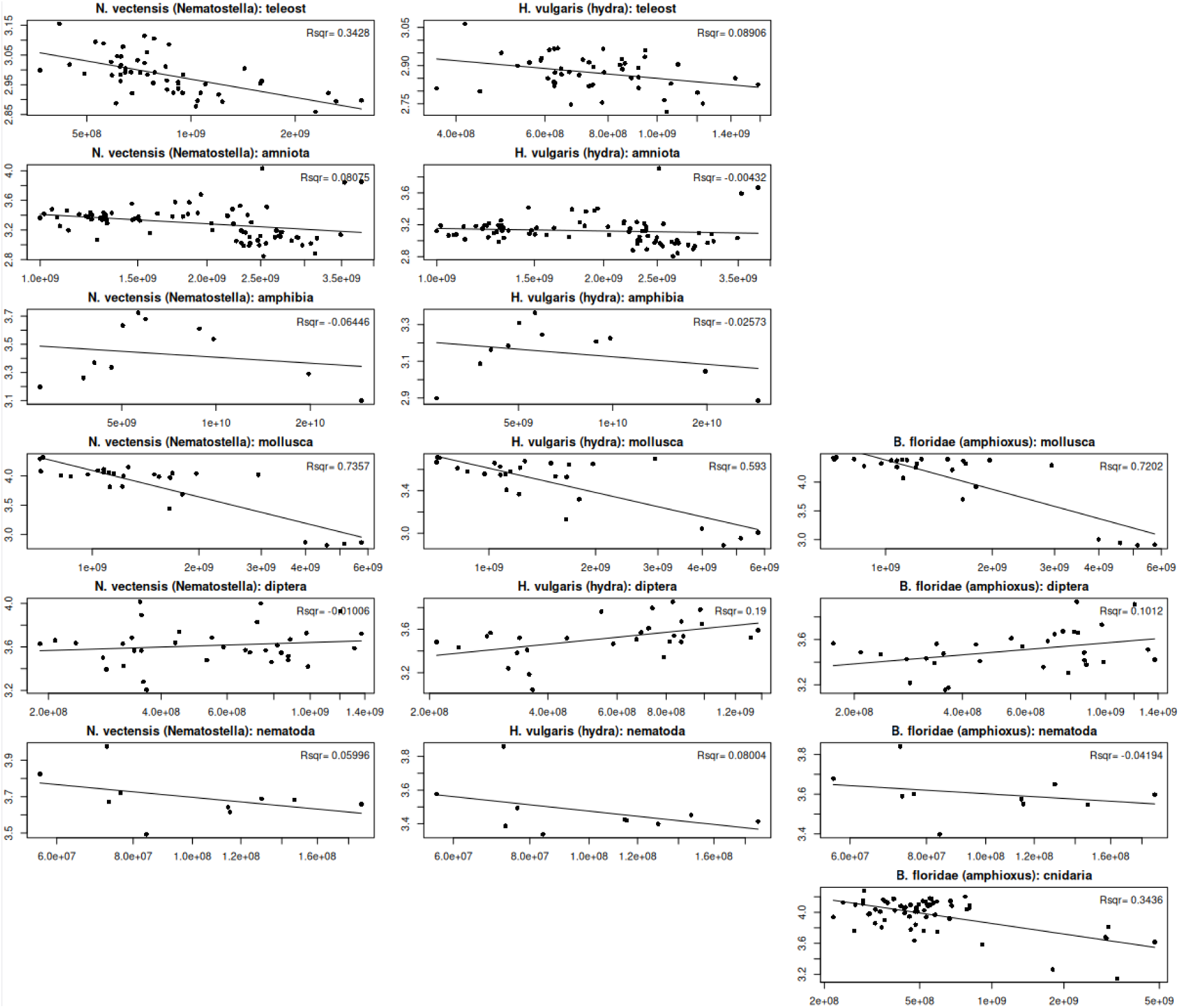
Mixing entropy (braiding entropy) tends to be inversely proportional to the genome size. Rows from top to bottom: *Nematostella vectensis* and *Hydra vulgaris* (respectively) pairwise comparisons to several clades (teleosts, tetrapods, molluscs, dipterans, nematodes), Branchiostoma floridae comparisons to molluscs, dipterans, nematodes and cnidarians. A regression line was fit to log(genome size). Braid entropy was computed with the braidlab package in MATLAB.

